# Protein hyperacylation links mitochondrial dysfunction with nuclear organization

**DOI:** 10.1101/2020.10.23.350892

**Authors:** John Smestad, Micah McCauley, Matthew Amato, Yuning Xiong, Juan Liu, Yi-Cheng Sin, Jake Ellingson, Yue Chen, Fatimah Al Khazal, Brandon Wilbanks, Jeong-Heon Lee, Tamas Ordog, Ioulia Rouzina, Mark Williams, Jason W. Locasale, L. James Maher

## Abstract

Cellular metabolism is linked to epigenetics, but the biophysical effects of metabolism on chromatin structure and implications for gene regulation remain largely unknown. Here, using a broken tricarboxylic acid (TCA) cycle and disrupted electron transport chain (ETC) exemplified by succinate dehydrogenase subunit C (SDHC) deficiency, we investigated the effects of metabolism on chromatin architecture over multiple distance scales [nucleosomes (∼10^2^ bp), topologically-associated domains (TADs; ∼10^5^ – 10^6^ bp), and chromatin compartments (10^6^ – 10^8^ bp)]. Metabolically-driven hyperacylation of histones led to weakened nucleosome positioning in multiple types of chromatin, and we further demonstrate that lysine acylation directly destabilizes histone octamer-DNA interactions. Hyperacylation of cohesin subunits correlated with decreased mobility on interphase chromatin and increased TAD boundary strength, suggesting that cohesin is metabolically regulated. Erosion of chromatin compartment distinctions reveals metabolic regulation of chromatin liquid-liquid phase separation. The TCA cycle and ETC thus modulate chromatin structure over multiple distance scales.

## Introduction

Mammalian epigenomic states are tightly coupled to cellular metabolism through multiple mechanisms (Dai et al., 2020; Etchegaray and Mostoslavsky, 2016; Reid et al., 2017), but an understanding of the effects of metabolic perturbations on the hierarchical structure of chromatin is in its very early stages. For example, pioneering studies examining direct effects of metabolism-driven epigenetic alterations on chromatin structure demonstrated that oncometabolite inhibition of TET1/2 DNA demethylases relevant to several human malignancies (Caramazza et al., 2010; Letouze et al., 2013; Toro et al., 2003; Yan et al., 2009) results in accumulation of DNA hypermethylation preventing the binding of CCCTC-binding factor (CTCF) to genomic DNA, resulting in loss of insulation between topologically-associated domains (TADs) and aberrant gene expression (Flavahan et al., 2019; Flavahan et al., 2016). There is also growing recognition that defects in the mitochondrial tricarboxylic acid (TCA) cycle and electron transport chain (ETC) are common in diabetes, heart disease, and cancer (Le et al., 2019; Miranda-Goncalves et al., 2018; Rosario et al., 2018; Tzika et al., 2018). In each of these cases, genetic and environmental contributions are hypothesized to drive metabolic alterations with corresponding epigenetic perturbations. Chromatin disorganization has been described in each of these diseases (Bysani et al., 2019; Corces et al., 2018; Costantino et al., 2017; Kleppe et al., 2018; Rosa-Garrido et al., 2017), though analyses infrequently assess the full hierarchical range of chromatin structure. It is unclear whether metabolically-driven epigenetic perturbations cause the chromatin structure changes in these diseases.

Here we demonstrate that mitochondrial TCA cycle and ETC dysfunction, exemplified through genetic knockout (KO) of succinate dehydrogenase subunit C (SDHC) in immortalized mouse embryonic fibroblasts (iMEFs), results in disruption of chromatin architecture at multiple distance scales including nucleosomes (∼10^2^ bp), topologically-associated domains (TADs; ∼10^5^ – 10^6^ bp), and chromatin compartments (10^6^ – 10^8^ bp). Further, we show that hyperacylation of nucleosome and non-nucleosome chromatin proteins is responsible for this coupling between metabolism and the epigenome. At the distance scale of nucleosomes, we demonstrate that SDHC KO results in a nucleosome lysine hyper-pan-acylation phenotype correlating with increased inter-dyad distances and weakened positioning of nucleosomes along the linear genome. These effects are partially reversible by treatment with histone acetyltransferase inhibitors (HATi). Our studies of direct effects of acyl PTMs on the biophysical properties of nucleosomes demonstrate that bulk nucleosomal acylation weakens histone-DNA interactions, enhancing nucleosome breathing, and promoting histone octamer diffusion along DNA molecules with reconstituted nucleosomes in the absence of externally-applied force or nucleosome remodeling activity. This suggests that histone hyperacylation and thermal energy are sufficient to drive nucleosome repositioning along DNA. Through profiling of genome-wide topological contact maps, we detect enhanced TAD boundary strength in SDHC KO cells, correlating with enhanced stability of cohesin complexes on chromatin in fluorescence recovery after photo-bleaching (FRAP) experiments and with hyperacylation of multiple cohesin subunits identified through targeted SILAC proteomic analysis. These results suggest that the stability of cohesin on chromatin may be metabolically regulated. At the distance scale of chromatin compartments, we show that SDHC KO results in decreased compartmentalization, preferentially impacting heterochromatic B compartments, and correlating with an equilibration of epigenomic marks between A/B compartments. We verify this phenomenon of eroded chromatin compartmentalization upon SDHC KO using immunofluorescence confocal microscopy and volumetric analysis of single cells, with partial rescue of the phenotype observed by treatment with HATi. We additionally demonstrate that a similar phenomenon of attenuated chromatin compartmentalization is observed upon pharmacological inhibition of ETC complex V with oligomycin A, an effect similarly rescued by treatment with HATi. Finally, we demonstrate that pharmacological inhibition of sirtuin 1 (SIRT1) by EX-527 is sufficient to induce attenuated chromatin compartmentalization, directly implicating SIRT1 in this process. Thus, perturbation of TCA cycle and ETC metabolism, exemplified by SDHC KO, modulates chromatin structure at three distance scales by acylation-dependent effects on nucleosome and non-nucleosome chromatin proteins.

## Results

### TCA cycle disruption causes a protein pan-hyperacylation phenotype

To study effects of TCA cycle and ETC dysfunction, we leveraged a tissue culture model involving normal and SDHC-knockout iMEFs (Figure S1A) (Smestad et al., 2018a; Smestad et al., 2018b; Smestad and Maher, 2019). Using these cells, we first assessed impacts of SDHC KO on cellular metabolism using untargeted metabolomic analysis. This revealed statistically-significant perturbation of >200 polar metabolites (Dataset S1). Chemical similarity enrichment analysis using ChemRICH (Barupal and Fiehn, 2017) revealed strong increases in several metabolite structural classes, including amino acids, dipeptides, and, interestingly, carnitines (Figure 1A). Broad increases were noted for short-, medium-, and long-chain acyl-carnitines, as well as reduction in levels of free L-carnitine (Figure 1B). These changes were associated with defective mitochondrial ultrastructure (Figure S1B) and reduction of ETC complex subunits (Smestad et al., 2018a). This correlation is consistent with prior reports linking ETC dysfunction with increased fatty acid oxidation (FAO) (Chen et al., 2018; Vazquez et al., 2015) and accumulation of acyl-carnitines (Audano et al., 2019), and may be related to the known physical association of FAO enzymes with ETC complexes (Wang et al., 2010; Wang et al., 2019b). Acyl-carnitine concentrations correspond to the concentrations of their respective reactive acyl-CoAs (Liu et al., 2015). The latter are capable of protein acylation at lysine residues (Simithy et al., 2017), suggesting to us the possibility that protein hyperacylation may be a consequence of SDHC KO.

**Figure 1.**
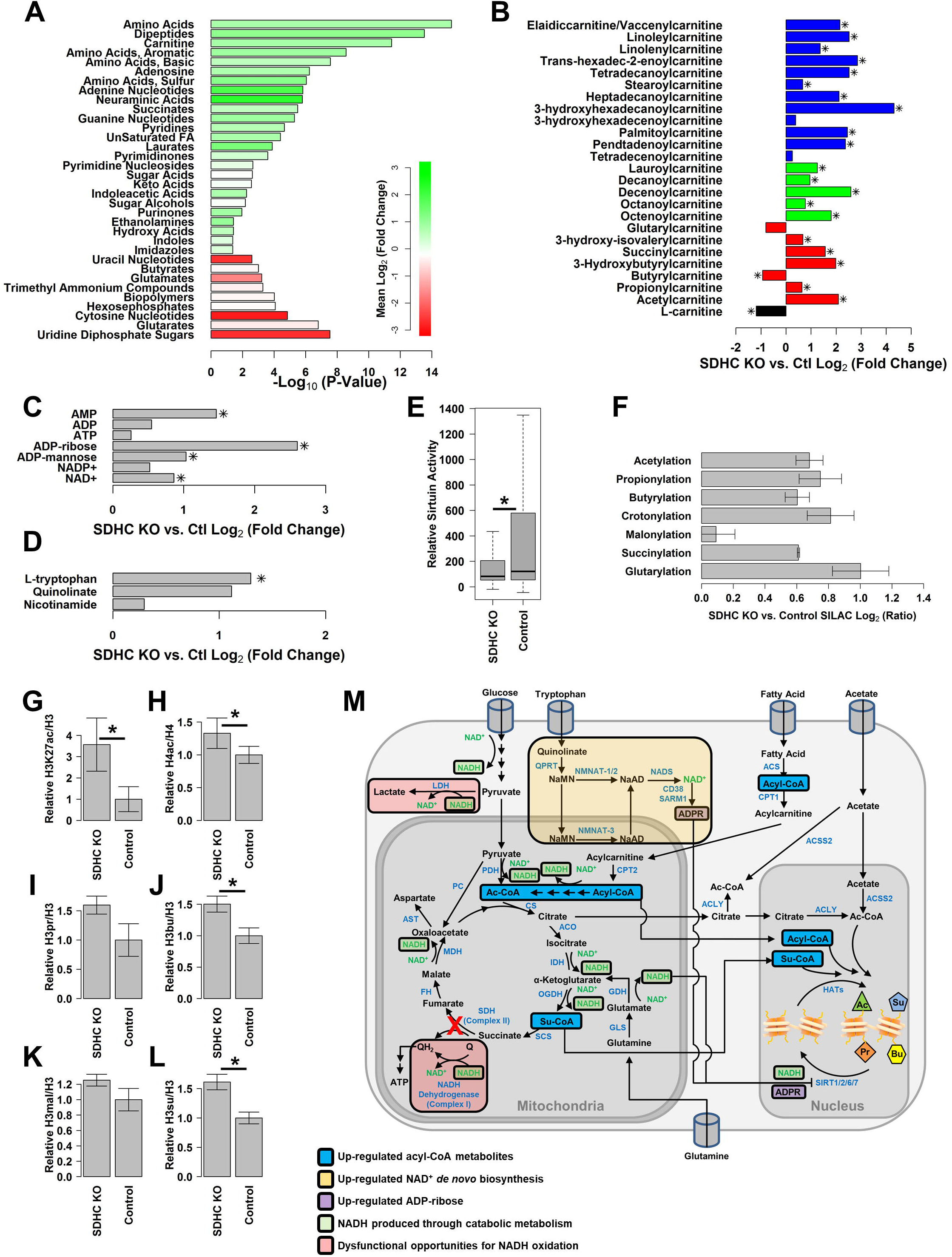
SDHC KO causes a pan-hyperacylation phenotype affecting chromatin. (A) ChemRICH structural similarity enrichment analysis of metabolic perturbations detected in SDHC KO cells relative to control cells. Colors indicate mean log_2_(fold change) for the respective metabolite structural classes. (B) Changes in acyl-carnitine metabolites detected in SDHC KO cells relative to control cells (* denotes p-value < 0.05 by two-sided heteroscedastic t-test, adjusted by FDR for multiple comparisons). Bar color denotes acyl chain length [red: short (< 7), green: medium (7 - 12), blue: long (> 12)]. (C) Changes in adenine nucleotides detected in SDHC KO cells relative to control cells. Statistical analysis was as in (B). (D) Changes in NAD^+^ biosynthetic precursors detected in SDHC KO cells relative to control cells. Statistical analysis was as in (B). (E) Quantification of relative sirtuin activity in SDHC KO and control cells by flow cytometry using an acylation-dependent, genetically-encoded fluorescent probe (* denotes p-value < 0.05 by Wilcox rank-sum test). (F) SILAC proteomics aggregate site-specific quantification ratios of acylated peptides normalized by corresponding protein abundance in SDHC KO and control cells. (G) Western blot quantification of H3K27ac normalized to total H3 (* denotes p-value < 0.05 by two-sided t-test). (H) Western blot quantification of H4ac (K5, K8, K12, and K16) normalized to total H4 (* denotes p-value < 0.05 by two-sided t-test). (I) Western blot quantification of pan-propionyl H3 normalized to total H3. (J) Western blot quantification of pan-butyryl H3 normalized to total H3 (* denotes p-value < 0.05 by two-sided t-test). (K) Western blot quantification of pan-malonyl H3 normalized to total H3. (L) Western blot quantification of pan-succinyl H3 normalized to total H3 (* denotes p-value < 0.05 by two-sided t-test). (M) Schematic illustration of metabolic effects observed in SDHC KO cells contributing to the chromatin hyperacylation phenotype. Error bars indicate standard deviations in all cases.

Our analysis of enriched metabolite structural classes additionally revealed a striking increase in adenine nucleotides (Figure 1A), with notable inclusion of NAD^+^ and related metabolite ADP-ribose (Figure 1C), correlating with increases in tryptophan and quinolinate (precursors in *de novo* NAD^+^ biosynthesis), but not nicotinamide (a precursor in the NAD^+^ salvage pathway) (Hassa et al., 2006; Stein and Imai, 2012) (Figure 1D). Endogenous production of ADP-ribose from NAD^+^, known to occur via multiple mechanisms (Hassa et al., 2006), is correlated with increased expression NAD^+^ glycohydrolases (Lee and Zhao, 2019) in the absence of other notable change in expression of NAD^+^-consuming and ADP-ribose generating enzyme activities (Figure S1C). ADP-ribose has previously been described as an endogenous sirtuin inhibitor (Madsen et al., 2016), suggesting to us the possibility of sirtuin inhibition in SDHC KO cells. Levels of nicotinamide, another known endogenous sirtuin inhibitor, were not changed upon SDHC loss (Figure 1D). Independent measurement showed the NAD^+^/NADH ratio to be decreased upon SDHC KO (Figure S1D), correlating with reduced levels of the protein subunits of NADH Dehydrogenase (ETC complex I) and an increase in lactate dehydrogenase (LDH) (Figure S1E), as well as increased lactate and decreased pyruvate (Figure S1F). These effects suggest that SDHC KO cells unsuccessfully induce LDH to rescue the abnormally low NAD^+^/NADH ratio, further supported by observations of the synthetic lethality of LDH loss with SDH loss (Bancos et al., 2013; Smestad et al., 2018b). While recent work has suggested that physiological NADH concentrations do not inhibit relevant sirtuins (Madsen et al., 2016), the collective metabolomic changes suggested to us that sirtuin activities may be inhibited upon SDHC KO.

Because of the relevance of sirtuin activity to protein acylation levels, we assessed endogenous sirtuin activity in SDHC KO and control cells using a genetically-encoded fluorescent reporter system. This system is sensitive to activities of SIRT1, SIRT2, SIRT3, and SIRT5 (Xuan et al., 2017). Monitoring by flow cytometry provided detection of sirtuin activity in SDHC KO and control cells (Figure S1G). Indeed, comparison of sirtuin activities (eGFP fluorescence) in the relevant subset of dual mCherry^+^ and eGFP^+^ cells created from SDHC KO and control lines revealed a significant decrease in sirtuin activity in SDHC KO cells relative to control (Wilcox rank sum p-value: 0.01, Figure 1E). This change did not correlate with alteration of sirtuin protein expression or subcellular localization (Figure S1H).

The combination of elevated concentrations of reactive acyl-CoAs and sirtuin inhibition is predicted to synergistically increase lysine PTM acylation marks on cellular proteins including nucleosomes (Li et al., 2016; Simithy et al., 2017; Vaquero et al., 2004; Vaquero et al., 2006). We therefore leveraged previously-reported SILAC proteomic data for SDHC KO and control cell lines (Smestad et al., 2018a) to quantitate lysine acylation in high-abundance peptides. This analysis revealed profound elevation of multiple lysine acyl PTM marks in SDHC KO cells, including acetylation, propionylation, butyrylation, crotonylation, succinylation, and glutarylation (Figure 1F). To examine whether hyperacylation included histone proteins, we performed Western blotting of nuclear extracts using antibodies specific to various representative histone acetylation marks including H3K27ac and H4K(5, 8, 12, 16)ac as well as antibodies specific for propionyllysine, butyryllysine, malonyllysine, and succinyllysine, with signal normalization to total histone. This analysis revealed H3K27 and H4K(5, 8, 12, 16) hyperacetylation, and increased H3 butyrylation and succinylation in the SDHC KO cell line relative to control (Figure 1G-L). These data support a model in which TCA cycle and ETC chain dysfunction, exemplified by SDHC KO, elevate multiple acyl PTMs, with effects extending to histone proteins in the nucleus (Figure 1M).

### Histone hyperacylation upon SDHC KO alters nucleosome positioning

Having demonstrated that SDHC KO results in a protein hyper-pan-acylation phenotype affecting nucleosomes, we next used this model system to ask if dysfunction of the TCA cycle and ETC influences nucleosome positioning along the linear genome. Histone acetylation has previously been proposed to alter nucleosome stability and positioning by histone lysine charge neutralization causing attenuation of favorable electrostatic interactions with DNA (Fenley et al., 2010; Fenley et al., 2018), and between histone tails (Dion et al., 2005; Otterstrom et al., 2019; Zhang et al., 2017a), as well as recruitment of ATP-dependent chromatin remodelers (Clapier et al., 2017). The mechanisms by which other acylation marks may impact chromatin structure are less clear, but a similar model invoking core histone lysine charge neutralization has been proposed for propionylation, butyrylation, and crotonylation (Fenley et al., 2018). To study the combined effects of nucleosome pan-hyperacylation in SDHC KO cells, we first performed automated annotation of epigenomic states across the genome using CUT&RUN and ChIP-seq data from SDHC KO and control cell lines (Figure S2A-H) and the ChromHMM algorithm (Ernst and Kellis, 2017) to generate a 12-state epigenomic annotation (Figure S2I-N). We then performed ATAC-seq in SDHC KO and control cells (Dataset S2), followed by an analysis of nucleosome positions using the NucleoATAC analysis pipeline (Schep et al., 2015), examining the impact of SDHC KO on nucleosome positioning within 10-Mbp random samples of genomic regions mapping to each of the epigenomic states determined by ChromHMM. Unsurprisingly, ATAC-seq coverage was enriched for euchromatic chromatin states (Figure S2O). Differences in nucleosomal inter-dyad distances between SDHC KO and control cells were determined for each of the identified epigenomic states. This revealed that SDHC KO cells have statistically-significant increases in inter-dyad distances at euchromatic chromatin regions including CTCF binding sites, simple repeats, enhancers, and gene promoters (Figure 2A). Differences were also calculated for nucleosome fuzziness scores (a metric of nucleosome positioning at the called position) along the linear genome, revealing weakened positioning upon SDHC KO for all chromatin epigenomic states except promoter downstream regions (Figure 2B). Analysis of ATAC-seq peaks additionally demonstrated a gain in total peak number for SDHC KO cells relative to control (Figure S3A-C). These changes are consistent with a model in which histone hyperacylation in SDHC KO cells results in attenuated nucleosome stability and weakened positioning.

**Figure 2.**
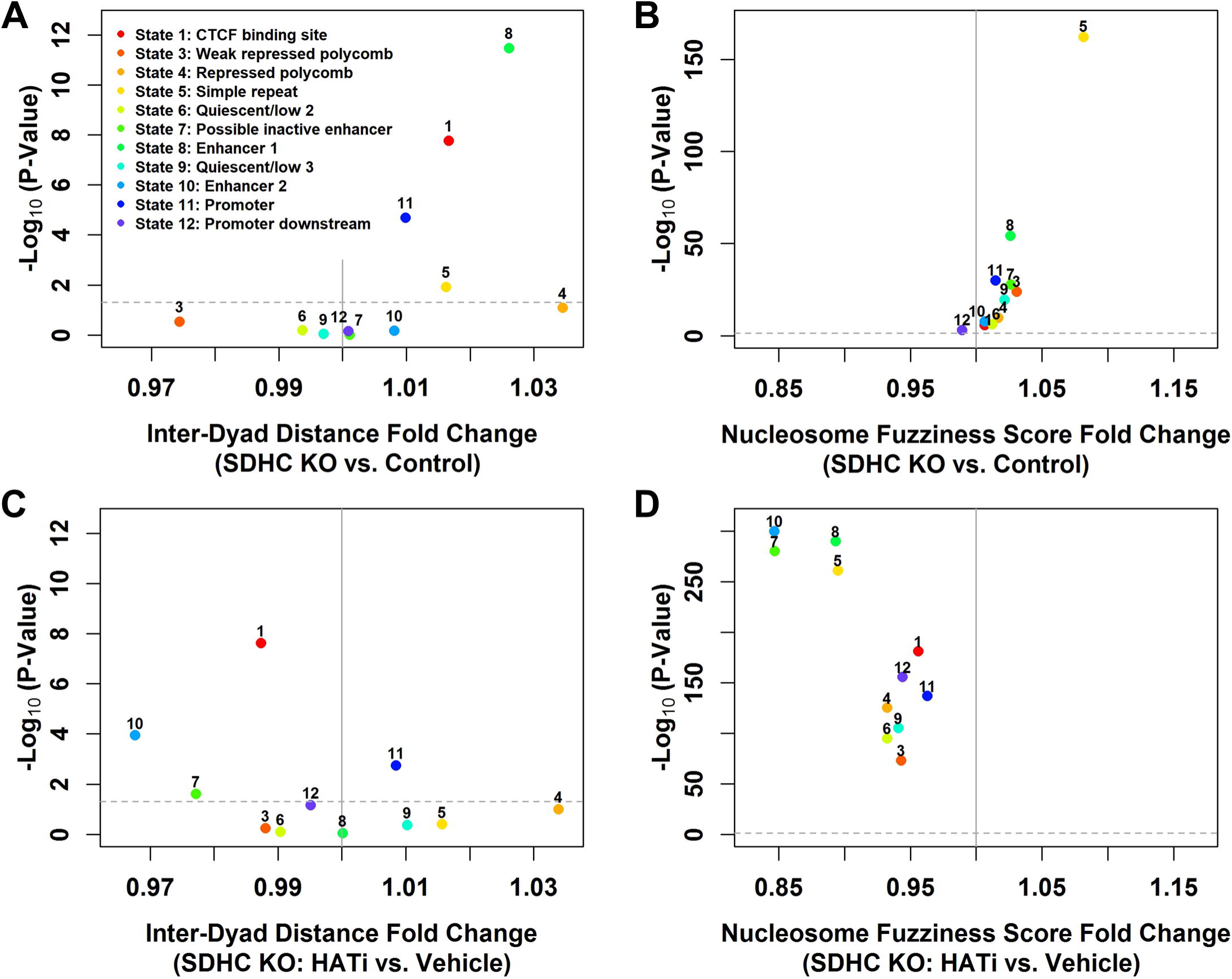
SDHC KO causes chromatin decompaction partially reversible by histone acetyltransferase inhibition. NucleoATAC analysis of nucleosome position and fuzziness (delocalization) was performed for ChromHMM-called chromatin states. X-axes denote parameter fold-change and y-axes denote –log_10_(p-value) calculated by Wilcox rank-sum test. (A) Analysis of nucleosome inter-dyad distance changes in SDHC KO cells relative to control. (B) Analysis of nucleosome fuzziness changes in SDHC KO cells relative to control. (C) Analysis of nucleosome inter-dyad distance changes in HATi-treated [MB-3 (100 µM) and C646 (10 µM) for 72 h] SDHC KO cells relative to vehicle-treated SDHC KO cells. (D) Analysis of nucleosome fuzziness changes in HATi-treated [MB-3 (100 µM) and C646 (10 µM) for 72 h] SDHC KO cells relative to vehicle-treated SDHC KO cells.

We next assessed whether weakened nucleosome positioning induced in SDHC KO cells could be reversed by chemical inhibition of histone acetylation. To this end, SDHC KO cells were treated with inhibitors of p300/CBP and Gcn5 histone acetyltransferases (10 µM C646 and 100 µM MB-3, respectively), resulting in ∼40% reduction in H3K27ac after 72 hours (Figure S3D,E). We then performed ATAC-seq profiling of SDHC KO cells treated with these histone acetyltransferase inhibitors (HATi) or vehicle, assessing impacts on nucleosome inter-dyad distances and fuzziness scores. Remarkably, HATi treatment of SDHC KO cells partially rescued the nucleosome inter-dyad distance distribution, and completely rescued the weakened nucleosome positioning for all chromatin regions assessed (Figure 2C,D). Similarly, ATAC-seq analysis revealed a decrease in total peak number for SDHC KO cells treated with HATi relative to vehicle (Figure S3F-H). These results demonstrate that weakened nucleosome positioning upon SDHC KO can be reversed through targeted reduction of histone acetylation by HATi, implicating histone acetylation as a functionally-important acylation mark controlling nucleosome positioning *in vivo*.

### Histone hyperacylation weakens nucleosomes *in vitro*

To assess direct biophysical effects of nucleosome acylation, including acylation marks that are difficult to individually manipulate *in vivo*, we developed a single-molecule *in vitro* assay. We performed optical tweezers experiments after treatment of reconstituted nucleosome arrays (Figure 3A) with different acyl-CoAs to quantify non-enzymatic acylation impacts on nucleosome stability. Under increasing tension, DNA has been previously shown to separate from the octamer in two distinct phases. At stretching forces up to 5 pN, the outer ½ turns are gradually disrupted as contacts between the DNA and the histone tails of the (H3-H4)_2_ tetramer are displaced along with partial disruption of interactions with the H2A/H2B core dimer (Figure 3B). Direct interactions between the inner DNA loop and histone helices in the tetramer and the H2A/H2B dimers (referred to as ‘strong sites’) are abruptly and individually disrupted at stretching forces above 10 pN (Brower-Toland et al., 2005a; Brower-Toland et al., 2002; McCauley et al., 2019). Figure 3C shows typical force-extension data for a control array and for arrays treated with acetyl-CoA and succinyl-CoA.

**Figure 3.**
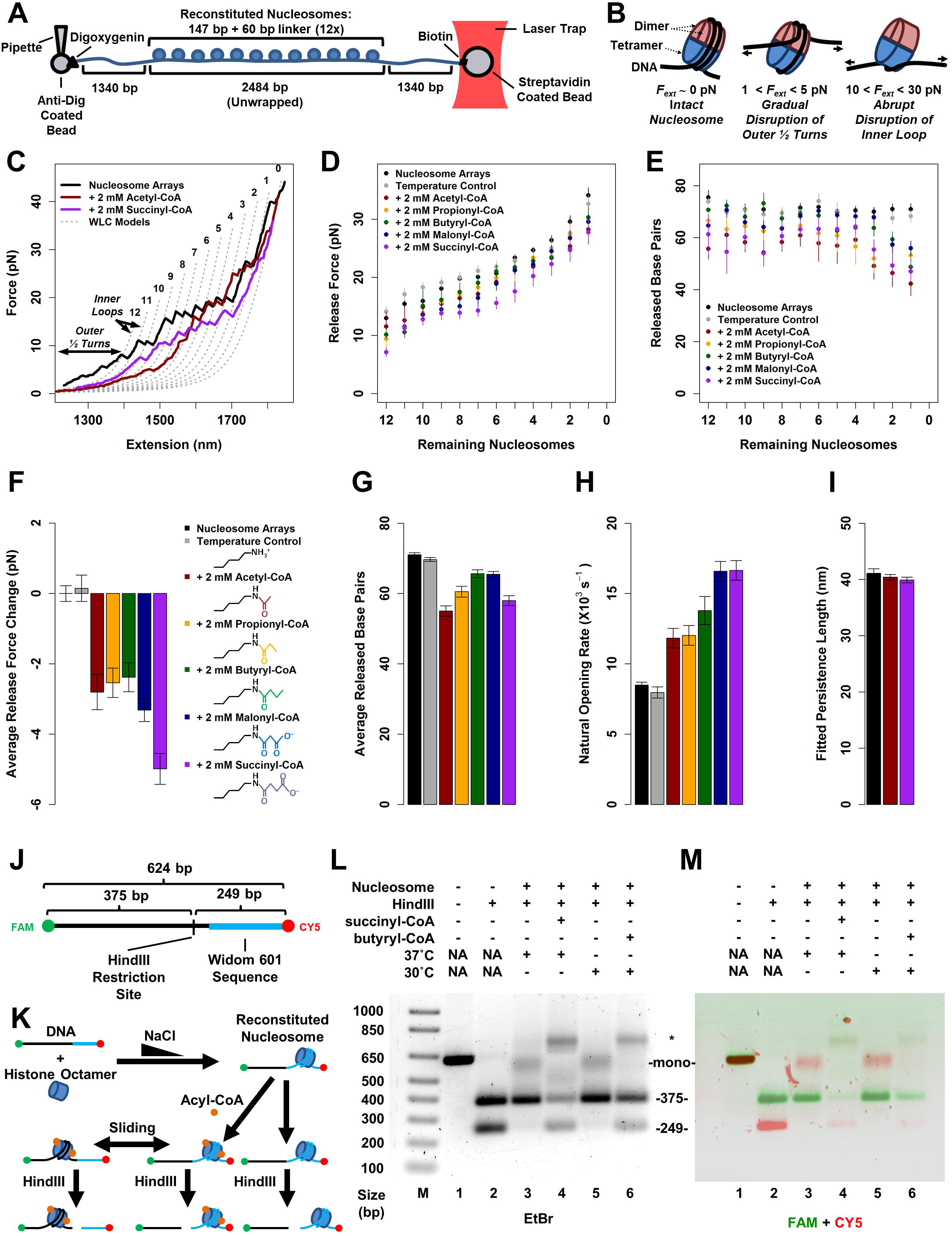
Acylation weakens nucleosomes *in vitro*. (A) Reconstituted nucleosome array analysis using optical tweezers. (B) Schematic of two stages of force-induced DNA release; the two outer half turns of DNA release below 5 pN while the inner loop of DNA is disrupted at higher forces. (C) Plot of force vs. extension analyzing an array of 12 nucleosomes reconstituted on linearized plasmid DNA disrupted under increasing tension (black). Colors indicate pre-incubation of arrays with 2 mM acyl-CoAs (red: acetyl-CoA, purple: succinyl-CoA). Gray traces indicate fits to the worm-like chain polymer model, each separated by 78-bp contour lengths, as described in the supplemental methods. (D) Measured release force for each nucleosome within an array for untreated arrays without incubation (black, *n* = 16), arrays incubated at room temperature without addition of acyl-CoAs (grey, *n* = 10), and arrays treated with acetyl-CoA (red, *n* = 8), propionyl-CoA (yellow, *n* = 8), butyryl-CoA (green, *n* = 12), malonyl-CoA (blue, *n* = 15), and succinyl-CoA (violet, *n* = 8). (E) DNA length released upon destabilization of each nucleosome measured at the force shown in D. (F) Averaged release force change per nucleosome for untreated arrays vs. treated arrays that result in the lysine modifications shown in the key. (G) Averaged released base pairs per nucleosome for untreated and treated arrays. (H) Fitted natural rate of DNA-nucleosome opening. (I) Fitted persistence lengths of DNA exposed to acetyl-CoA (red) or succinyl-CoA (violet). Error bars indicate standard error in the mean for the respective measurements. Color codes in panels G-I are as in panel F. (J) Schematic representation of DNA construct used for nucleosome reconstitutions. (K) Illustration of experimental workflow used for gel analysis of acylation-dependent effects on nucleosome migration. (L) Ethidium-stained 1.5% agarose gel showing nucleosome reconstitution (lanes 3-6) followed by the indicated acylation (or mock) treatment and HindIII digestion. (M) Same as L but overlay of false-color images reflecting fluorescent detection with FAM and Cy5 filters (green and red), respectively. DNA fragment sizes (249 bp, 375 bp) are shown along with position of mononucleosome at Widom 601 sequence (“mono”) and co-migrating novel nucleosome species (*) with both FAM and Cy5 signals.

The behavior of reconstituted nucleosome arrays acylated by exposure to acyl-CoAs in aqueous buffer differs from untreated arrays in three critical ways. First, the unmodified array shows a force-extension plateau below 5 pN of stretching force, an effect entirely absent in the treated arrays, indicating that histone acylation essentially eliminates stabilizing interactions between the exit/entry DNA ½ turns and the histones (Figure 3C). Thus, all subsequent data characterize the disruption of the inner DNA loop from the strong sites. These strong site interactions, characterized by ‘rips’ in the extension data, are disrupted by lower stretching forces in treated arrays (Figure 3D), while the amount of DNA that remains wrapped also decreases, indicating weakened histone – DNA interactions for each nucleosome in the treated arrays (Figure 3E). Averaged over all arrays, we see that lysine modification by acyl groups of increasing length and negative charge is increasingly destabilizing (Figure 3F,G). Fitting this disruption data determines the natural nucleosome opening (or ‘breathing’) rate in the absence of force (Figure 3H). This opening rate increases for all histone acylations, growing with the length of the added group, and nearly doubling for the longest chains that also reverse the net lysine charge from a cation to an anion. As expected, the persistence length of DNA was unchanged by acyl-CoA treatment (Figure 3I), indicating that acylation of histone lysines does not modulate the intrinsic stiffness of DNA itself.

The finding that nucleosomes are weakened by acylation, which increases nucleosomal opening, was further supported by the results of ensemble *in vitro* transcription experiments utilizing DNA template sequences bearing single reconstituted nucleosomes, demonstrating decreased transcriptional pausing by an elongating bacteriophage RNA polymerase at acylated nucleosomes (Figure S4A,B). This result suggests that acylated nucleosomes pose a lower barrier to the passage of translocating molecular machines (Chang et al., 2013; Studitsky et al., 1995). Together, these *in vitro* data support the conclusion that chemical acylation weakens nucleosomes, reducing disruption force, extent of DNA wrapping, and stability to RNA polymerase elongation, with all acylation marks displaying qualitatively similar effects that increase in magnitude with the negative charge on the acylation mark.

Since our ATAC-seq data demonstrated weakened nucleosome positioning along the linear genome of SDHC KO cells, and because *in vitro* acylation of histone lysines weakened histone octamer interactions with DNA, we next assessed whether histone acylation can promote nucleosome sliding within a linear DNA construct. We reconstituted single histone octamers onto a 624-bp duplex DNA construct containing a Widom 601 nucleosome-positioning sequence (Figure 3J) (Lowary and Widom, 1998). Reconstituted nucleosomes were then exposed to acyl-CoAs in aqueous buffer, followed by restriction digestion to produce DNA fragments amenable to nucleosome positioning analysis by agarose gel electrophoresis (Figure 3K). In the absence of acyl-CoA treatment, reconstituted nucleosomes remain localized exclusively to the restriction fragment containing the Widom 601 nucleosome positioning sequence. Remarkably, acylation causes nucleosome migration, resulting in repositioned nucleosomes along the DNA construct, including nucleosomes localized on the restriction fragment lacking the positioning sequence (Figure 3L,M; Figure S4C). This acylation-induced nucleosome migration is driven only by thermal energy, independent of any external force or ATP-dependent nucleosome remodeling complex, demonstrating that multiple types of lysine acylation marks promote nucleosome sliding along linear DNA.

### SDHC KO increases TAD boundary strength and cohesin stability on chromatin

We next examined the effects of TCA cycle and ETC dysfunction due to SDHC KO on higher levels of chromatin organization, beginning with topologically-associated domains (TADs). For this purpose, we generated three-dimensional chromatin contact maps for SDHC KO and control cells using a modified Hi-C method (eHi-C) (Lu et al., 2018). We leveraged the Juicer analysis pipeline (Durand et al., 2016) and the Knight-Ruiz method for normalization of chromatin contact matrices (Knight and Ruiz, 2013). Since cell cycle stage is a known determinant of chromatin topological organization (Nagano et al., 2017), we first ruled out differences in cell cycle distribution between SDHC KO and control cell populations using DAPI/Geminin staining and analytical flow cytometry to show similar cell proportions in G1, early S, and late S-G2 phases (Figure S5A,B). Analysis of biological replicate eHi-C experiments for SDHC KO and control cells using HiCRep (Yang et al., 2017) revealed good agreement between replicates, with larger differences observed between SDHC KO and control groups (Figure 4A). Insulation scores for all chromosomes were then calculated for each biological replicate using a 10-Kbp resolution contact matrix and a sliding 500 Kbp × 500 Kbp window (Figure S5C), as previously described (Crane et al., 2015). Analysis of inter-replicate insulation score correlations further demonstrated good agreement between biological replicates, with larger differences observed between SDHC KO and control groups (Figure S5D). TAD boundaries were called jointly for SDHC KO and control datasets using the insulation score method (Crane et al., 2015), which is sensitive for identifying TADs across a wide range of read depths (Zufferey et al., 2018). This identified 2286 TADs with boundary strength >0.5, allowing us to assess differences in TAD boundary strength between SDHC KO and control cell lines. Remarkably, this analysis revealed a profound enhancement in TAD boundary strength in SDHC KO cells relative to controls (Figure 4B-E). The notion that TAD boundary strength can be linked to TCA cycle and ETC dysfunction provides evidence that TCA cycle activity can alter genome organization by a mechanism known to be important for gene regulation.

**Figure 4.**
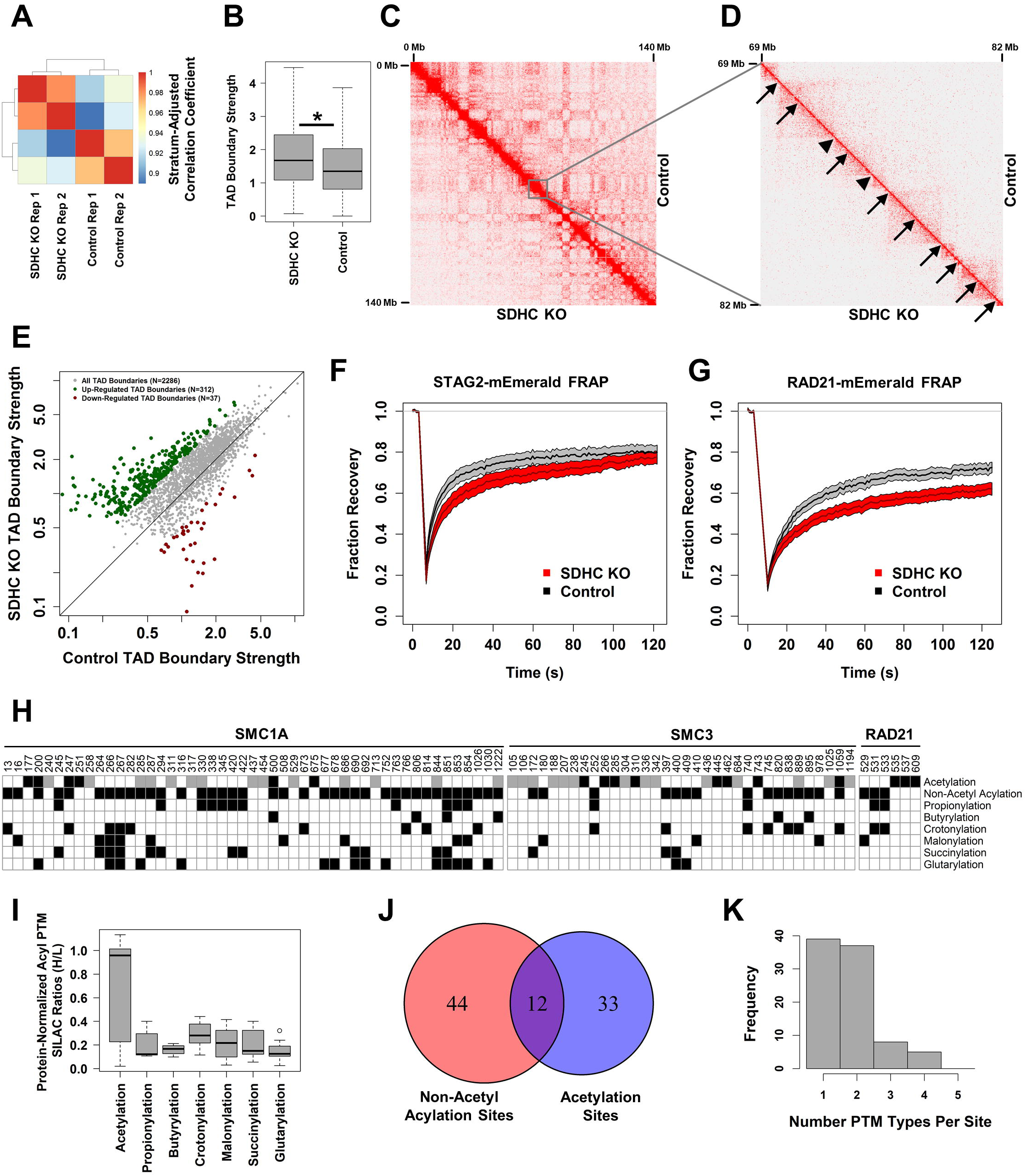
SDHC KO results in enhanced TAD boundary strength and increased cohesin stability on chromatin, correlating with cohesin hyperacylation. (A) HiCRep stratum-adjusted correlation coefficients calculated at 1 Mbp resolution for biological replicate eHi-C datasets. (B) TAD boundary strengths in SDHC KO and control cells (* denotes p-value < 0.05 by Wilcox rank-sum test). (C) KR-normalized eHi-C contact matrices for SDHC KO (x-axis) and control (y-axis) eH-C datasets showing chromosome 3 (0-140 Mbp) at 250 Kbp resolution. (D) Subset of (C) showing SDHC KO (x-axis) and control (y-axis) eH-C chromosome 4 (69-82 Mbp) at 25 Kbp resolution. Arrows indicate positions of identified TAD boundaries. Arrows with stems indicate increased boundary strength in SDHC KO cells relative to control [log_2_(fold-change) > 1]. Arrowheads without stems indicate TADs with unchanged boundary strength. (E) Plot of TAD boundary strength in SDHC KO and control cells. Colors indicate TADs with increased [log_2_(fold-change) > 1[and decreased [log_2_(fold-change) < −1] boundary strength in SDHC KO relative to control cells. (F) Aggregate normalized FRAP curves for mEmerald-STAG2 transiently expressed in SDHC KO (N=12) and control (N=13) cells. Ranges indicate SEM. (G) Aggregate normalized FRAP curves for mEmerald-RAD21 transiently expressed in SDHC KO (N=24) and control (N=15) cells. Ranges indicate SEM. (H) Acyl PTMs identified on cohesin subunits via SILAC proteomics. Amino acid positions on respective cohesin subunits are indicated. Colors indicate PTM presence (gray: unchanged SDHC KO vs. control; black: uniquely identified in SDHC KO). (I) Estimated protein-normalized acyl PTM SILAC H/L ratios. For PTMs only identified in SDHC KO line, H/L ratio estimated as 1/(S/N ratio). (J) Venn diagram of acetylation and non-acetyl acylation site locations. (K) Histogram of number of PTM types per site identified.

Recent evidence suggests that TADs are dynamic structures, forming and dissolving during interphase on minute timescales as a function of cohesin dissociation from chromatin (Hansen et al., 2018; Hansen et al., 2017). We therefore designed experiments to test if enhanced TAD stability in SDHC KO cells could be attributed to altered stability of cohesin on interphase chromatin. We performed fluorescence recovery after photobleaching (FRAP) experiments leveraging transient expression of mEmerald-labeled cohesin components STAG2 and RAD21 in SDHC KO and control cells. Intensities of pre-bleach regions of interest (ROIs) in these experiments were similar between SDHC KO and control cells (Figure S5E,F). Importantly, analysis of FRAP profiles revealed increased stability of both STAG2 and RAD21 cohesin components on chromatin in SDHC KO cells relative to controls (Figure 4F,G, Movies S1-4), consistent with the increase in TAD boundary strength observed in bulk chromatin contact maps.

To understand the basis for increased stability of cohesin on chromatin in SDHC KO cells, we checked for differences in abundance of CTCF, cohesin subunits, or known cohesin regulators as detected by SILAC proteomics. The abundance of these components did not significantly differ between SDHC KO and control cell lines (Figure S5G), suggesting that the differences in TAD boundary strength are not due to altered expression of cohesin subunits or known cohesin regulators. We also monitored CTCF site occupancy from ATAC-seq footprint analysis, revealing similar occupancy profiles in both SDHC KO and control cell lines (Figure S4H,I). Because DNA binding by CTCF is known to be inhibited by cytosine methylation at CpG sites (Hashimoto et al., 2017), and a previous report has linked SDH loss to DNA hypermethylation with inhibition of CTCF binding (Flavahan et al., 2019), we checked the DNA methylation status of CTCF CpG sites using reduced representation bisulfite sequencing (RRBS). This study revealed no significant difference in CTCF CpG methylation between SDHC KO and control cells (Fig S5J). Together, these data suggest that genomic occupancy by CTCF does not differ between SDHC KO and control cells. Post-translational acetylation of cohesin subunit SMC3 by ESCO1/ESCO2 has been reported to modulate cohesin binding to interphase chromatin (Kawasumi et al., 2017). However, no differences in SMC3 K105/K106 acetylation were observed between SDHC KO and control cells (Figure S5K,L).

In the absence of alterations in known determinants of cohesin stability on interphase chromatin, we studied the possibility that post-translational modification of cohesion subunits differs between SDHC KO and controls cells. We therefore performed SILAC proteomics in SDHC KO and control cell lines. Following SILAC labeling of control cells using heavy (^13^C_6_-labeled) lysine and arginine, and culture of SDHC KO cells in equivalent media containing light (^12^C_6_-labeled) lysine and arginine, cohesin complexes were purified from cell lysates by immunoprecipitation and gel electrophoresis prior to proteomic analysis of pooled heavy and light (H+L) peptides. Analysis of raw peptide abundance H/L ratios ranged from 1.5-2.3, indicating an excess of control (heavy) peptides relative to SDHC KO (light) peptides. Analysis of acyl PTMs identified a total of 129 acyl PTMs at 89 unique sites on SMC1A, SMC3, and RAD21. Strikingly, the vast majority of identified acyl PTMs (101/129), including all non-acetyl acylations, were detectable only in SDHC KO cells, despite the raw peptide abundance favoring the control specimens (Figure 4H). Conservative estimation of SILAC H/L ratio from measured peptide intensities and quantification of site-specific signal-to-noise ratios, normalized to input peptide abundance, suggests a dramatic increase in non-acetyl acylation PTMs on cohesin subunits in SDHC KO (Figure 4I). We also found that locations of acetylation PTMs and non-acetyl acylation PTMs are generally non-overlapping, suggesting different regulatory mechanisms (Figure 4J). In addition, the majority of identified sites bear either 1 or 2 acyl PTMs, with a small subset displaying 3 or 4 different acyl PTMs (Figure 4K). This suggests the possibility of site-specific competition. These observations are consistent with a model in which dysfunction of the TCA cycle and ETC results in pan-hyperacylation of cohesin subunits, correlating with altered cohesin biophysics on chromatin.

### Chromatin hyperacylation upon SDHC KO reduces chromatin compartmentalization

Finally, we examined effects of TCA cycle and ETC dysfunction on chromatin compartmentalization, leveraging eHi-C datasets generated for SDHC KO and control cells. Strikingly, SDHC KO results in attenuated chromatin compartmentalization (Figure 5A). Analysis of compartment eigenvector autocorrelations revealed that attenuated compartmentalization is concentrated in B (heterochromatic) compartments relative to A (euchromatic) compartments (Figure 5B-E). Compartment identity was found to be generally conserved, with 92% of compartments having the same identity between SDHC KO and control cell lines. We checked whether decreased chromatin compartmentalization correlated with altered distribution of epigenetic marks, considering the portion of the genome with conserved compartment identity between SDHC KO and control cell lines. This analysis identified a remarkable erosion of compartment-specific epigenetic marks, resulting in a trend toward equilibration of epigenetic states (Figure 5F-H) that correlates with compartment structural erosion observed in autocorrelation analysis. Specifically, we see that levels of A compartment marks, including H3K27ac and H3K4me3, are reduced in A compartments and increased in B compartments upon SDHC KO (Figure 5F,G). The B compartment mark, H3K27me3, exhibits a similar pattern reflecting a net gain in A compartments and loss in B compartments (Figure 5H). Collectively, these data demonstrate that the observed weakening of chromatin compartmentalization upon TCA cycle and ETC dysfunction correlates with a dramatic perturbation of epigenetic states that blurs the distinction between compartment identities.

**Figure 5.**
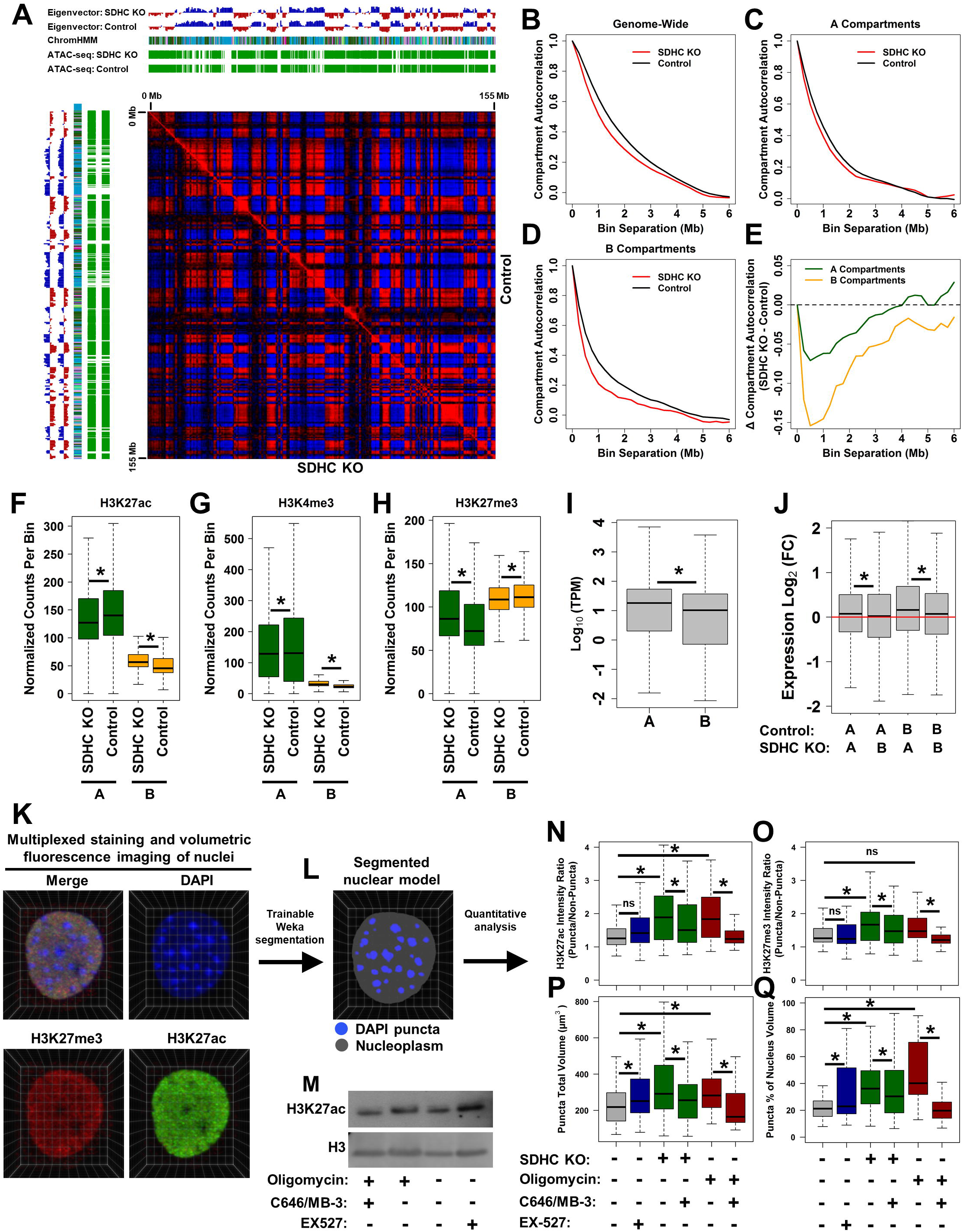
SDHC KO causes weakened chromatin compartmentalization, rescued by inhibition of histone acetyltransferases. (A) Pearson correlation coefficient matrix calculated from eHi-C contact matrices of SDHC KO (x-axis) and control (y-axis) chromosome 4 (0-155 Mb) at 500 Kbp resolution. Compartment eigenvectors, ChromHMM annotations, and ATAC-seq peaks are shown. (B) Genome-wide compartment autocorrelations calculated as a function of bin separation along the linear genome for SDHC KO and control cells. (C) Compartment autocorrelations within “A” compartments. (D) Compartment autocorrelations within “B” compartments. (E) Calculated differences in compartment autocorrelation between SDHC KO and control cells. (F-H) Boxplots of CPM-normalized reads per 250 Kbp bin for genomic regions mapping to A and B compartments with conserved identities in SDHC KO and control cells for (F) H3K27ac CUT&RUN, (G) H3K4me3 CUT&RUN, and (H) H3K27me3 ChIP-seq. (I) Boxplot of RNA-seq TPM-normalized expression quantities for transcripts mapping to A and B compartments in control cells. (J) Analysis of RNA-seq TPM-normalized expression changes for genes mapping to compartments that convert between A/B in SDHC KO cells relative to genes mapping to compartments that do not convert. (K) Representative images from multiplexed staining and volumetric imaging of fixed cell nuclei. Voxel dimension is 140 nm square horizontally and 320 nm vertically. (L) Representative segmented nuclear model derived from DAPI 3D image stack using Trainable Weka Segmentation. (M) Western blot of H3K27ac and total H3 for control cells treated with oligomycin (1 µM for 24 h), oligomycin and C646/MB-3 (10 µM C646, 100 µM MB-3 for 24 h), or EX-527 (20 µM for 24 h). (N) Analysis of H3K27ac intensity ratio for DAPI-stained puncta relative to non-puncta regions of nucleus. (O) Analysis of H3K27me3 intensity ratio for DAPI-stained puncta relative to non-puncta regions of nucleus. (P) Quantification of DAPI puncta total volume per cell. (Q) Quantification of DAPI puncta percentage of nucleus volume. * denotes p-value < 0.05 by Wilcox rank-sum test.

**Figure 6.**
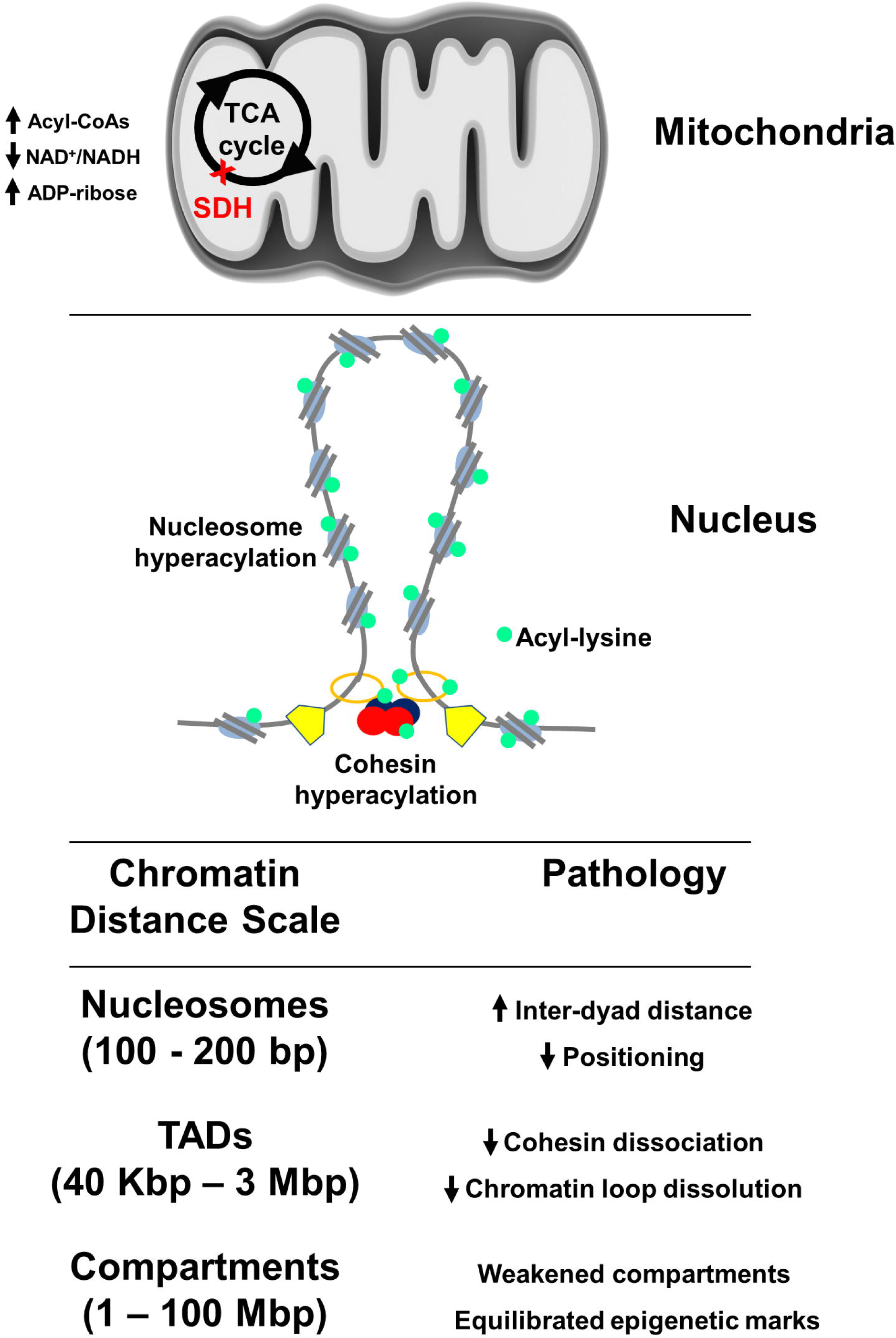
Schematic illustration of mechanisms by which mitochondrial electron transport chain dysfunction in SDHC KO affect chromatin structure.

Because chromatin compartment identity is a known determinant of transcriptional patterns (Lieberman-Aiden et al., 2009) (Figure 5I), we next assessed whether altered compartment structures correlate with transcriptional effects. Indeed, for the subset of compartments that flip A/B compartment state upon SDHC KO (∼8% of genome), we detect transcriptional changes that accompany this alteration in compartment identity (Figure 5J).

We verified the phenomenon of reduced chromatin compartmentalization using an orthogonal approach involving multiplexed immunostaining of fixed cells with antibodies specific to H3K27ac and H3K27me3. When followed by DAPI staining and high resolution confocal microscopy, this approach allows volumetric reconstruction of single nuclei to derive chromatin spatial information at the distance scale of chromatin compartments (Figure 5K), similar to a previously-reported method (Linhoff et al., 2015). Following image acquisition, Trainable Weka Segmentation (TWS) (Arganda-Carreras et al., 2017) was used to derive a pixel-based segmented nuclear model from DAPI fluorescence, classifying individual pixels as extra-nuclear, nucleoplasm, or DAPI heterochromatin puncta via a FastRandomForest algorithm comprised of 100 trees, each constructed from 7 random features trained on a total of 27,881 manually-annotated instances derived from images of SDHC KO and control cells (Figure 5L). 10-fold cross validation revealed the classifier to be robust, with 99.98% of manually-annotated instances correctly classified. This approach was applied to analyze SDHC KO and control cells, and cells exposed to ETC complex V inhibitor oligomycin A, and SIRT1 inhibitor EX-527, both of which increase histone H3K27 acetylation (Figure 5M). Effects of HATi in SDHC KO and oligomycin A-treated cells were also explored. Using CellProfiler (McQuin et al., 2018) for automated image analysis to assess H3K27ac and H3K27me3 puncta/nucleoplasm intensity immunofluorescence ratios for individual cells, we found that SDHC KO increases both H3K27ac and H3K27me3 marks in DAPI puncta relative to nucleoplasm, with reversal of these patterns upon HATi treatment (Figure 5N,O). A similar pattern of altered H3K27ac distribution was observed upon oligomycin A exposure, with corresponding rescue by co-treatment with HATi. A prior report has suggested that H3K27me3 marks a specific subset of heterochromatin not enriched in DAPI puncta (Linhoff et al., 2015). Accordingly, the observation of increased H3K27ac and H3K27me3 marks in DAPI heterochromatin puncta indicates reduced chromatin compartmentalization upon SDHC KO or oligomycin A exposure, with rescue of normal compartmentalization upon HATi treatment. Attenuated chromatin compartmentalization in TCA cycle and ETC dysfunction, with rescue by HATi treatment, was further supported by analysis of DAPI heterochromatin puncta volumes. Increased DAPI puncta volumes were found in both SDHC KO cells and oligomycin A-treated cells, again with rescue by HATi treatment (Figure 5P,Q). DAPI heterochromatin volumes were also increased upon EX-527 treatment, directly implicating SIRT1 in maintenance of chromatin compartmentalization, though an altered spatial distribution of H3K27ac and H3K27me3 was not observed. These results demonstrate that the TCA cycle and ETC regulate epigenetic marks modulating chromatin compartmentalization in living cells, consistent with recent evidence that chromatin liquid-liquid phase separation (LLPS) may be regulated by post-translational modification of histone proteins, including acetylation (Gibson et al., 2019b; Wang et al., 2019a).

## Discussion

### A protein hyperacylation phenotype is associated with dysfunction of the TCA cycle and ETC

Here we show that TCA and ETC dysfunction, triggered by SDHC loss, results in accumulation of acylcarnitines, NAD^+^ and ADP-ribose with a reduced NAD^+^/NADH ratio, sirtuin inhibition, and a protein pan-hyperacylation phenotype extending to histone and non-histone proteins in chromatin. We subsequently demonstrate acylation-dependent perturbations in chromatin structure over three discrete distance scales including mononucleosomes, TADs, and chromatin compartments. Epigenomic perturbation linked to TCA cycle and ETC dysfunction is relevant to both rare human diseases such as SDH-mutant paraganglioma and pheochromocytoma (Fishbein et al., 2017; Letouze et al., 2013), gastrointestinal stromal tumors (Gill et al., 2010; Gill et al., 2011), and IDH-mutant glioblastoma (Brennan et al., 2013), as well as common disorders including type II diabetes mellitus and heart failure. In the latter cases, ETC dysfunction with consequent sirtuin inhibition and a protein hyperacylation phenotype has been reported both for peripheral tissues in type II diabetes and cardiomyocytes in heart failure (Bagul et al., 2015; Lee et al., 2016). We predict that the epigenomic and chromatin structural changes we report upon SDHC loss will be mirrored in these disease states.

### Histone acylation weakens nucleosomes

We report ATAC-seq experiments that demonstrate increased nucleosome inter-dyad distances and de-localization upon SDHC KO, correlating with a histone lysine hyperacylation phenotype. These perturbations tend to be rescued by HAT inhibition, consistent with prior reports that nucleosome acetylation increases chromatin accessibility (Gorisch et al., 2005; Zhang et al., 2017b). Impacts of other acyl PTMs on nucleosome biophysical properties have previously been unclear, however. Fenley et al. (Fenley et al., 2018) proposed a core histone charge reduction model based on computational predictions for acylation marks on lysine residues near DNA including propionylation, butyrylation, and crotonylation. The majority of reported acyl-CoA modifications have been found within the tetrameric (H3-H4)_2_ tails (Anderson et al., 2001; Bintu et al., 2012; Brower-Toland et al., 2005b; Hebbes et al., 1994; Kebede et al., 2017; Simpson et al., 1978). Here we directly measure how total histone acylation affects the stability of individual nucleosomes, strikingly demonstrating that lysine modification by acyl-CoAs reduces nucleosome stability. Unwinding of the outer half turns of DNA from modified nucleosomes occurred spontaneously, consistent with weakened interactions between DNA and the preferentially acylated (H3-H4)_2_ tails, and explaining the known role of acylation in facilitating Pol II transcription (Bintu et al., 2012). We also quantified significant weakening of the off-dyad strong nucleosome sites anchoring the inner turn of DNA, detected as a reduction of the release stretching force and shrinking of the histone-DNA interaction site size, leading to a natural strong site opening rate increase by several fold. A comparable study of nucleosomal acylation catalyzed by the p300 enzyme also revealed decreased tetramer release forces with a modest reduction of DNA held by the tetramer tails, though a smaller number of lysine residues may have been modified (Brower-Toland et al., 2005a). We found that the effects of acylation progressively increase with the number of carbons of the added chain and upon change in lysine charge from +1 to −1. In the most extreme cases, chemical acylation destabilizes nucleosomes more than deletion of all histone tails. Finally, our fitted natural nucleosome opening rate, a reliable signature of the overall site stability, increases with longer (and more negatively charged) acyl modifications, suggesting an enhanced probability of histone octamer sliding along the DNA.

### Nucleosome acylation induces spontaneous migration along linear DNA

Currently recognized determinants of nucleosome positioning along the linear genome include DNA sequence composition, non-histone DNA binding proteins, ATP-dependent nucleosome remodelers, and the presence of transcription factors (Chereji and Clark, 2018). Our *in vitro* analysis strikingly demonstrates that nucleosome acylation is sufficient to promote repositioning, presumably by histone octamer sliding, along a linear DNA construct. This result shows that neutralizing or inverting lysine charges in histones lubricates the octamer-DNA interface, facilitating a fundamentally different kind of nucleosome mobility. The results of this experiment suggest that chromatin acylation *in vivo* may facilitate redistribution of nucleosomes along the linear genome. Testing this hypothesis *in vivo* is complicated by the challenge of targeted perturbation of individual acyl-CoAs without disrupting other metabolic pathways (Kustatscher et al., 2019).

### Hyperacylation of cohesin increases its stability on interphase chromatin

Cohesin mobility on interphase chromatin is known to be regulated by PTMs including phosphorylation (Bauerschmidt et al., 2011) and SMC3 K105/K106 acetylation (Kawasumi et al., 2017). Here we show that dysfunction of the TCA cycle and ETC results in decreased mobility of cohesin on interphase chromatin, and that this correlates with increased abundance of acyl PTMs on multiple cohesin subunits. Aside from SMC3 K105/106 acetylation, the vast majority of acetylation PTMs on cohesin subunits have unknown functions in regulating cohesin biophysics. In addition to acetylation, we here implicate non-acetyl acylations as having potential functional roles in cohesion regulation. Future studies utilizing *in vitro* systems will be needed to confirm that hyperacylation of cohesin, or introduction of site-specific lysine charge modifications, results in altered biophysics on chromatin. Assessment of site-specific contributions of individual acyl PTMs to cohesin stabilization on chromatin also remain to be determined. If future studies confirm that cohesin hyperacylation results in altered kinetics of chromatin binding, this would establish a direct mechanism by which formation of chromatin loops anchoring topological associated domains is metabolically regulated. Intriguingly, metabolic regulation of cohesin by sirtuins has previously been proposed, based both on analysis of functional overlap of sirtuin and cohesin genomic localization in yeast (Li et al., 2013), and by analysis of SIRT6 protein interaction networks in human cells (Simeoni et al., 2013). Modulation of cohesin function by metabolic effects has not previously been demonstrated. It remains unclear whether the non-acetyl acylation sites identified in our analysis are regulated by sirtuins, and further mechanistic work will be a high priority for future study.

### Chromatin hyperacylation weakens A/B compartmentalization

Links between epigenomic state and chromatin compartmentalization have been previously proposed (Jost et al., 2014; Nuebler et al., 2018). An intriguing hypothesis is that the separation of chromatin into A/B compartments is driven by chromatin-associated proteins that mediate the process of liquid-liquid phase separation (LLPS) (Larson et al., 2017; Strom et al., 2017). Evidence has also emerged that post-translational modifications of histone proteins may regulate chromatin phase separation through recruitment of chromatin-associated proteins that modulate LLPS (Wang et al., 2019a). A prior report has demonstrated that chromatin acetylation weakens LLPS *in vitro* (Gibson et al., 2019a), consistent with our observation that chromatin hyperacetylation in the context of TCA cycle and ETC dysfunction in living cells results in attenuated chromatin compartmentalization, and is rescued by HATi treatment. Here we extend the current paradigm by demonstrating a direct mitochondrial role in regulating chromatin A/B compartmentalization. It remains unclear whether the multiple dysregulated acyl PTMs in TCA cycle and ETC dysfunction share similar effects in perturbing chromatin compartmentalization. This will likely be best assessed by future *in vitro* studies examining the impact of individual nucleosome acylations on chromatin LLPS.

## Supporting information

Dataset S1. Polar metabolomics data

Dataset S2. ATAC-seq read mapping

Movie S1. FRAP mEmerald-RAD21 control

Movie S2. FRAP mEmerald-RAD21 SDHC KO

Movie S3. FRAP mEmerald-STAG2 control

Movie S4. FRAP mEmerald-STAG2 SDHC KO

## Author contributions

Conceptualization, J.S. and L.J.M; Methodology, J.S., L.J.M., Y.X., J.H.L., T.O., J.L., M.A., B.W., M.M., M.W., and Y.C.; Validation, J.S.; Formal Analysis, J.S., and Y.C.; Investigation, J.S., M.M., Y.X., M.A., F.K., B.W., J.L., Y.C.S., J.E., and Y.C..; Resources, L.J.M. and J.S.; Data Curation, J.S.; Writing – Original Draft, J.S.; Writing – Review & Editing, L.J.M., J.S., M.M. M.A., Y.X., J.L., Y.C.S., J.E., Y.C., F.A.K. B.W., J.H.L., T.O., J.W.L., M.W., and I.R.; Visualization, J.S.; Supervision, L.J.M., M.W., J.H.L., Y.C., J.W.L.; Project Administration, L.J.M.; Funding Acquisition, L.J.M, J.S.

## Acknowledgements

This work was facilitated by the Mayo Clinic Medical Genome Facility Sequencing Core, the Mayo Clinic Epigenetics Development Laboratory, the Mayo Clinic Microscopy and Cell Analysis Core Facility, the Mayo Clinic Research Computing Facility, and the University of Minnesota Proteomics Core Facility. We thank the staff of the Epigenomics Development Laboratory and Recharge Center (EDL) at Mayo Clinic for carrying out the ATAC-seq assays for this study. The EDL is supported in part by the Mayo Clinic Center for Individualized Medicine. We thank Peter Schultz (Scripps Research Institute) for the gift of pcDNA-mCherry-EGFP (K85TAG) and pCMVAcKRS-tRNAPyl plasmids used for sirtuin activity measurements. We thank Steven Henikoff (Fred Hutchinson Cancer Research Center, HHMI) for the gift of protein A-micrococcal nuclease fusion protein used for CUT&RUN profiling of epigenetic marks. We thank Karolin Luger (University of Colorado) for the gift of human histone octamers used for some nucleosome reconstitution experiments. We thank Georges Mer and Maria Victoria Botuyan (Mayo Clinic) for the gift of plasmid pET-15b-derived vectors for the expression of human histones H2A (pHISPP/H2A), H2B (pHISPP/H2B), H3.1 (pHISPP/H3.1), and H4 (pHISPP/H4), and technical assistance with other nucleosome reconstitution experiments. We thank Christina Murray for assistance with false-color image reconstruction in Fig. 3M.

## Supplemental figure titles and legends

**Figure S1.**
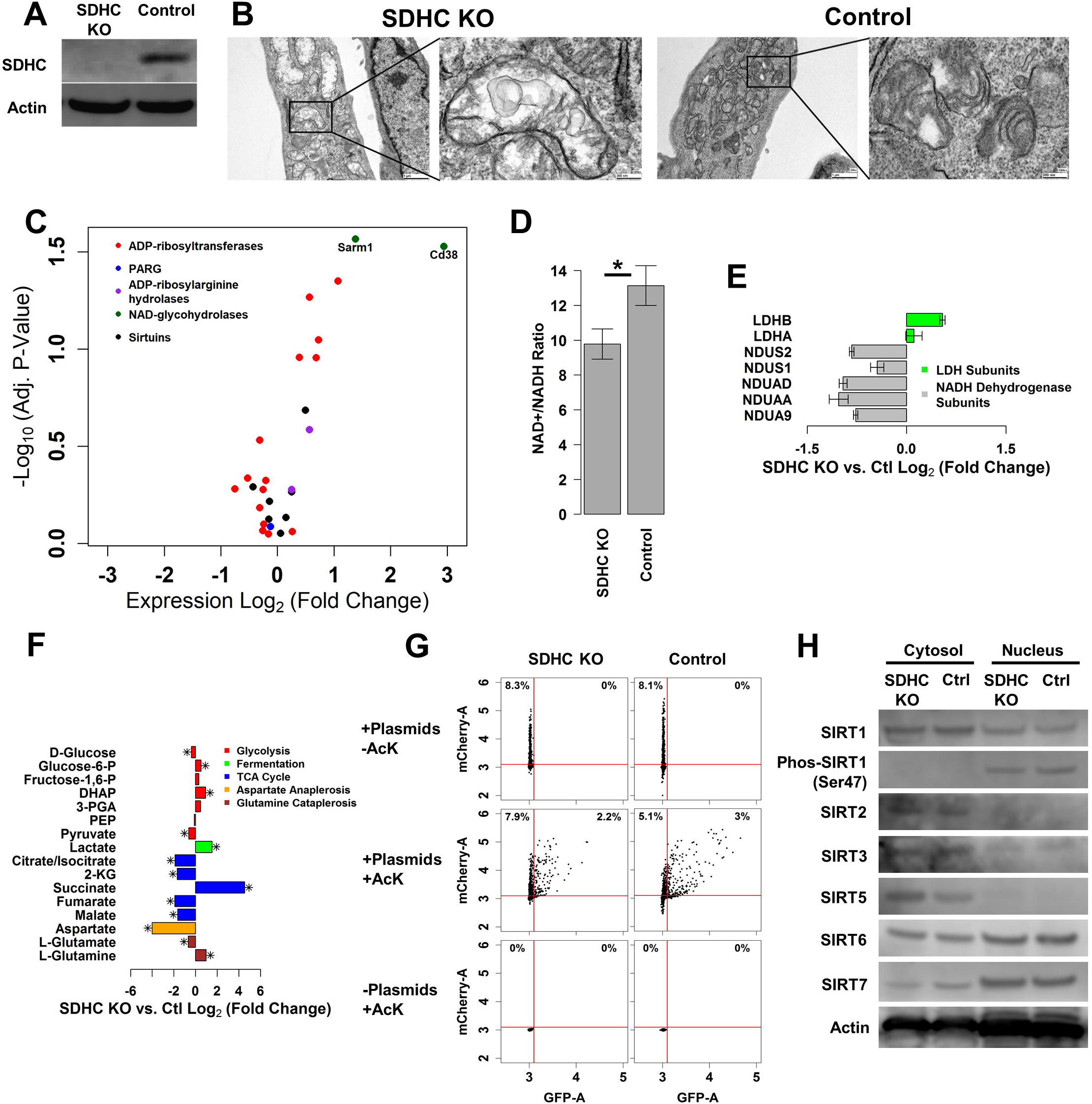
Metabolic analysis of SDHC KO and control cells, related to Figure 1. (A) Western blot of SDHC protein in SDHC KO and control cells. (B) Transmission electron microscopy analysis of mitochondrial morphology in SDHC KO and control cells. (C) RNA-seq analysis of ADP-ribose-producing biological activities in SDHC KO and control cells. (D) Quantification of NAD^+^/NADH ratio in SDHC KO and control cells (* denotes p-value < 0.05 by two-sided t-test). (E) SILAC proteomic quantification of LDH and NADH Dehydrogenase complex subunit expression change in SDHC KO cells relative to control. (F) Analysis of polar metabolites involved in glycolysis, fermentation, TCA cycle, aspartate anaplerosis, and glutamine cataplerosis in SDHC KO and control cells. (G) Flow cytometry analysis of sirtuin activity using genetically-encoded reporter. Conditions denoted “+ Plasmids” indicate transfection with pcDNA-mCherry-EGFP (K85TAG) and pCMVAcKRS-tRNAPyl plasmids. “+AcK” denotes the addition of acetyllysine to 5 mM concentration. (H) Western blot analysis of sirtuin protein levels in cytosolic and nuclear subcellular fractions for SDHC KO and control.

**Figure S2.**
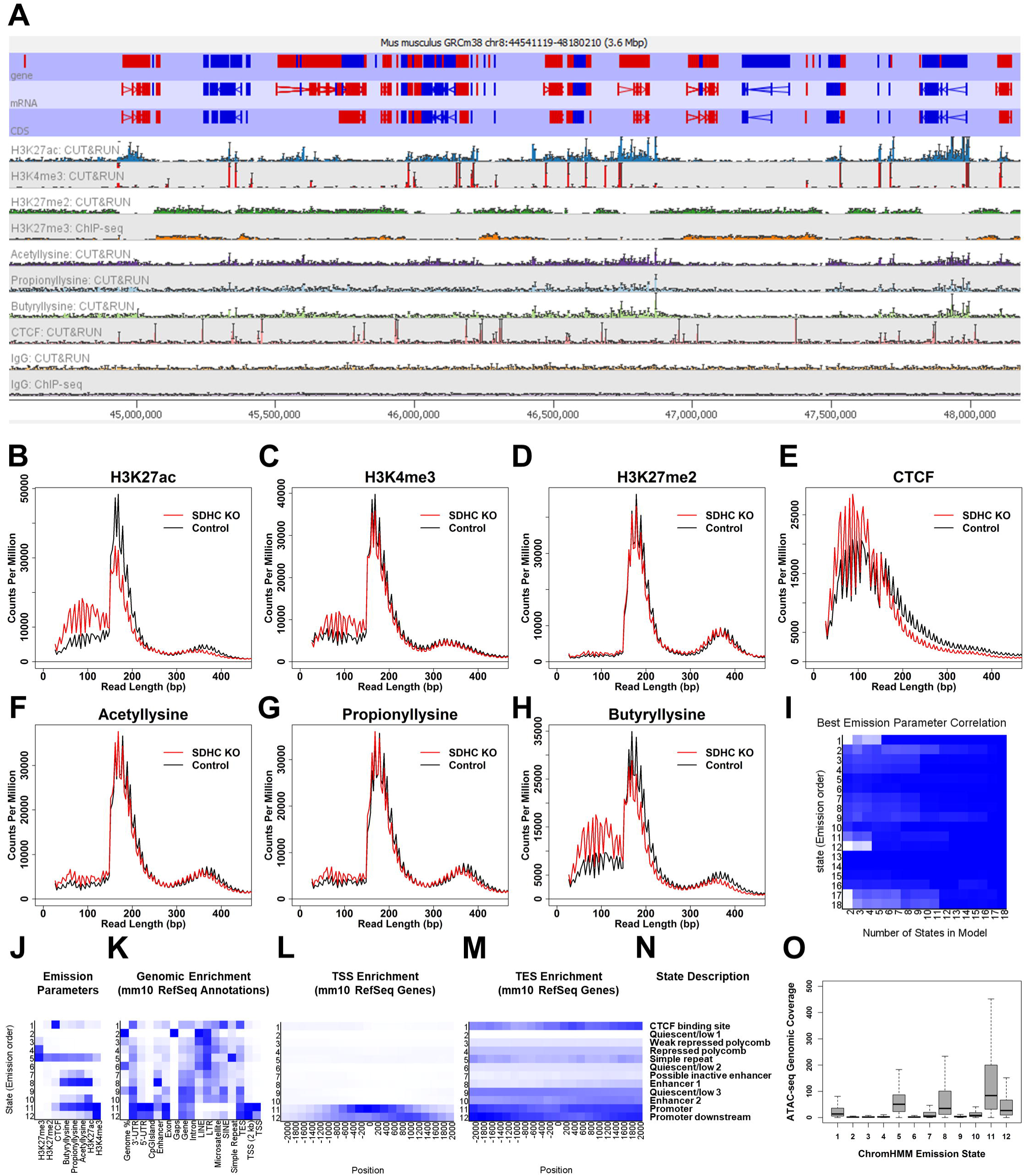
CUT&RUN analysis and generation of ChromHMM states, related to Figure 2. (A) SeqMonk genomic view of epigenomic profiles generated from CUT&RUN and ChIP-seq at 5 kbp resolution for control cell line. Error bars indicate STDEV for CPM-normalized probe quantities from biological replicate experiments. (B-H) Fragment length histograms from CUT&RUN profiling in SDHC KO and control cell lines using antibodies specific to H3K27ac, H3K4me3, H3K27me2, CTCF, acetyllysine, propionyllysine, and butyryllysine. (I) ChromHMM emission parameter correlations plotted as a function of number of states in stacked epigenomic model. (J) ChromHMM emission parameters. (K) ChromHMM genomic enrichments for RefSeq annotations. (L) ChromHMM TSS enrichment patterns. (M) ChromHMM TES enrichment patterns. (N) ChromHMM state descriptions. (O) Boxplot of ATAC-seq coverage for ChromHMM states.

**Figure S3.**
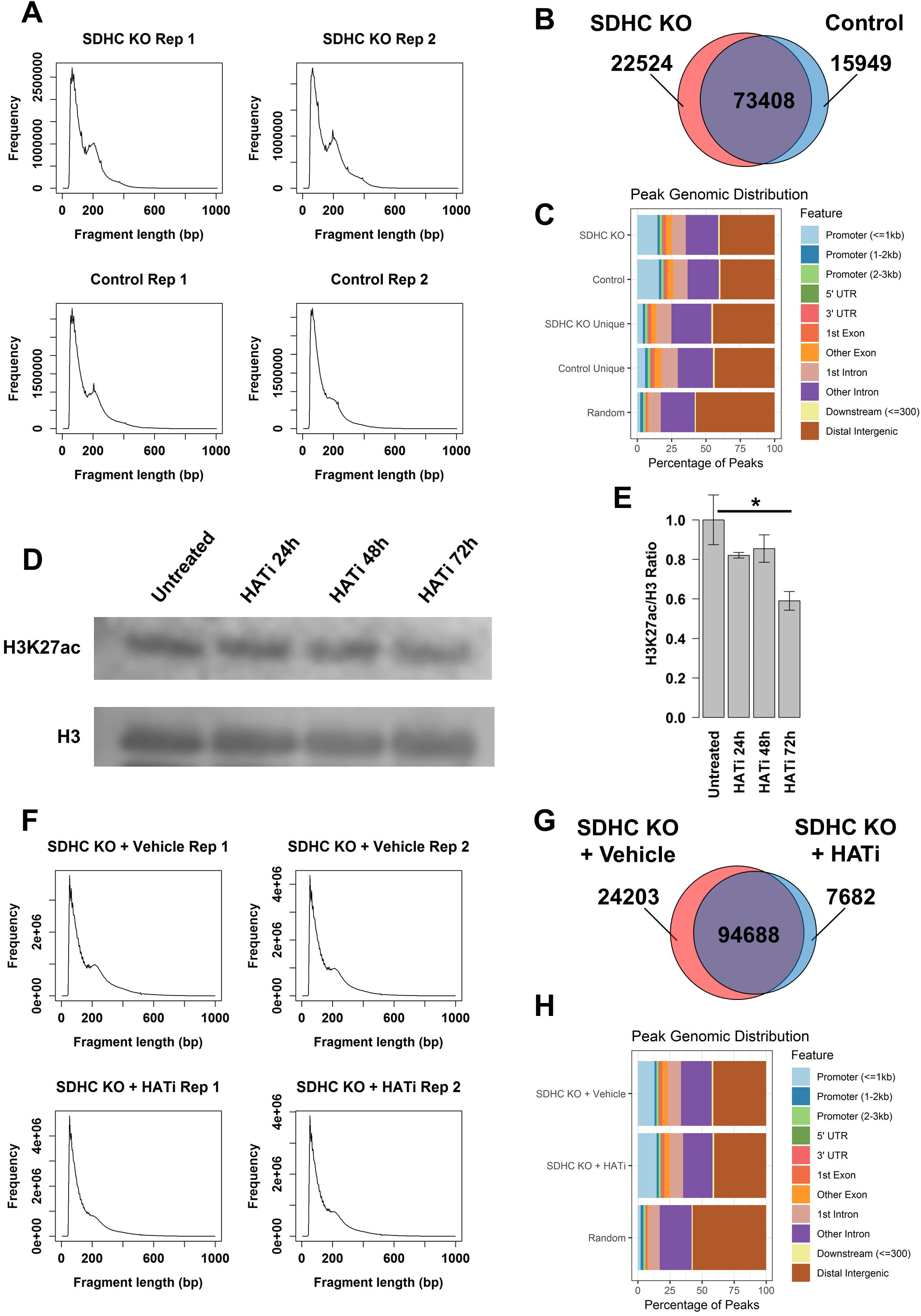
Supplemental ATAC-seq analysis and verification of SDHC KO H3K27ac rescue by HATi treatment, related to Figure 2. (A) Analysis of fragment length from ATAC-seq experiments in SDHC KO and control lines. (B) ATAC-seq peak numbers for SDHC KO and control lines identified by the MACS peak-calling algorithm. (C) Analysis of genomic localization of ATAC-seq peaks in SDHC KO and control cells. (D) Western blot analysis of H3K27ac and total H3 in SDHC KO cells treated with HATi (10 µM C646, 100 µM MB-3) for 24-72 h. (E) Quantification of relative H3K27ac/H3 ratio as a function of HATi treatment duration. * denotes two-tailed p-value < 0.05 by unpaired t-test. (F) Analysis of fragment length from ATAC-seq experiments in SDHC KO cells either treated with HATi (10 µM C646, 100 µM MB-3) or vehicle for 72 h. (G) ATAC-seq peak numbers for SDHC KO cells either treated with HATi (10 µM C646, 100 µM MB-3) or vehicle for 72 h identified by the MACS peak-calling algorithm. (H) Analysis of genomic localization of ATAC-seq peaks in SDHC KO cells either treated with HATi (10 µM C646, 100 µM MB-3) or vehicle for 72 h.

**Figure S4.**
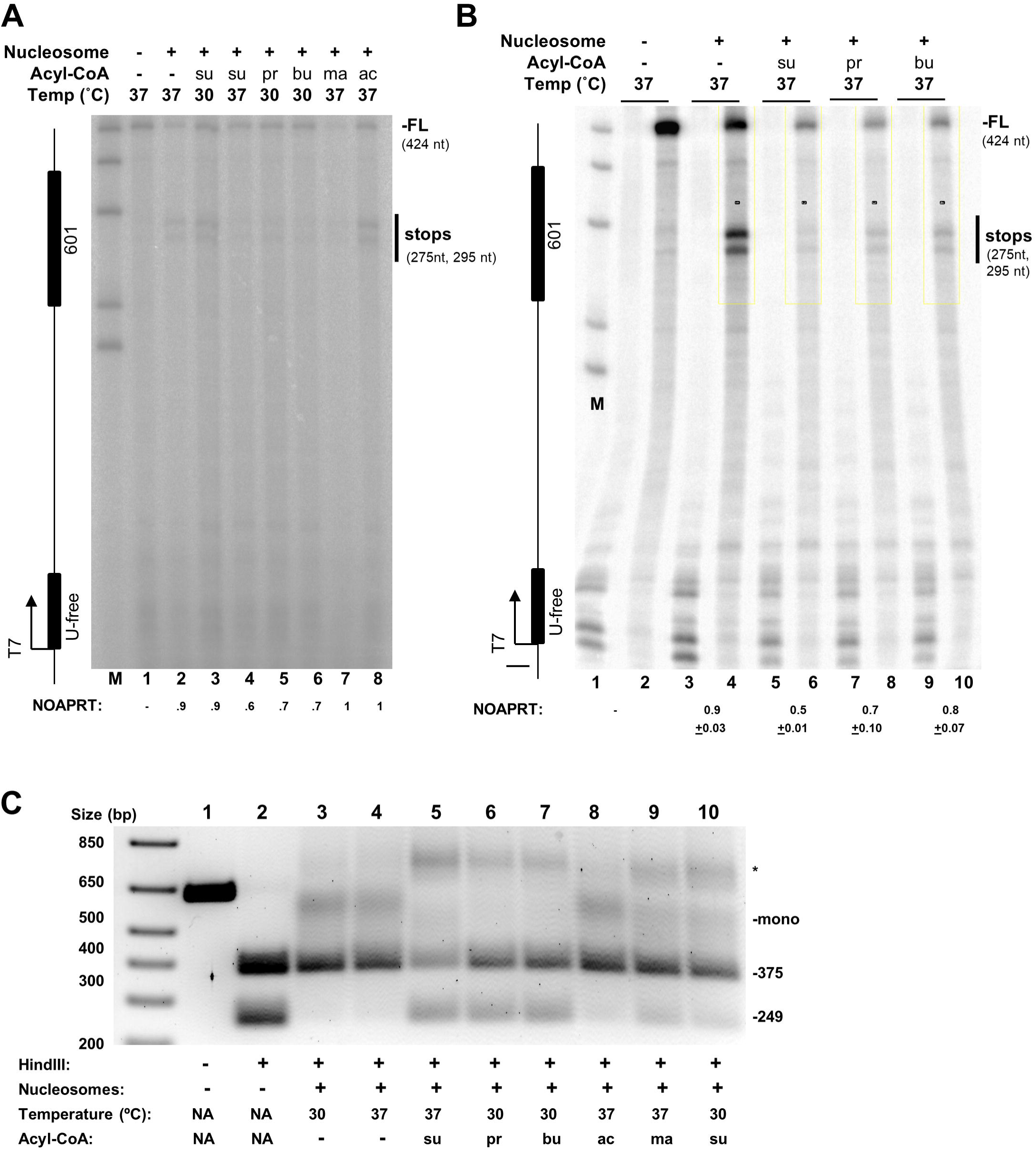
Supplemental analysis of nucleosome stability on reconstituted DNA constructs, related to Figure 3. (A,B) Analysis of nucleosome stability by monitoring T7 RNA polymerase transcription pausing and stopping during elongation at 4°C *in vitro* as described in methods. RNA transcripts were radiolabeled during preliminary open complex formation by transcription of a U-free cassette prior to cooling to 4°C and additional of all NTPs. Quantitation of two main stop sites near the center of the reconstituted nucleosome and normalization to the extent of template reconstitution (Nucleosome Obstacle Activity Per Reconstituted Template; NOAPRT) was as described in methods. Reduced NOAPRT values reflect less stable nucleosomes. *In vitro* nucleosome acylating agents and temperatures are indicated. (A) Example of results of single pilot study for multiple conditions. (B) Representative result and quantitation from independent studies (n = 3). (C) Agarose gel analysis of acylation-dependent effects on the stability of reconstituted nucleosomes, related to Figure 3K,L. Bands visualized with EtBr post-staining. Novel nucleosome species indicated as *.

**Figure S5.**
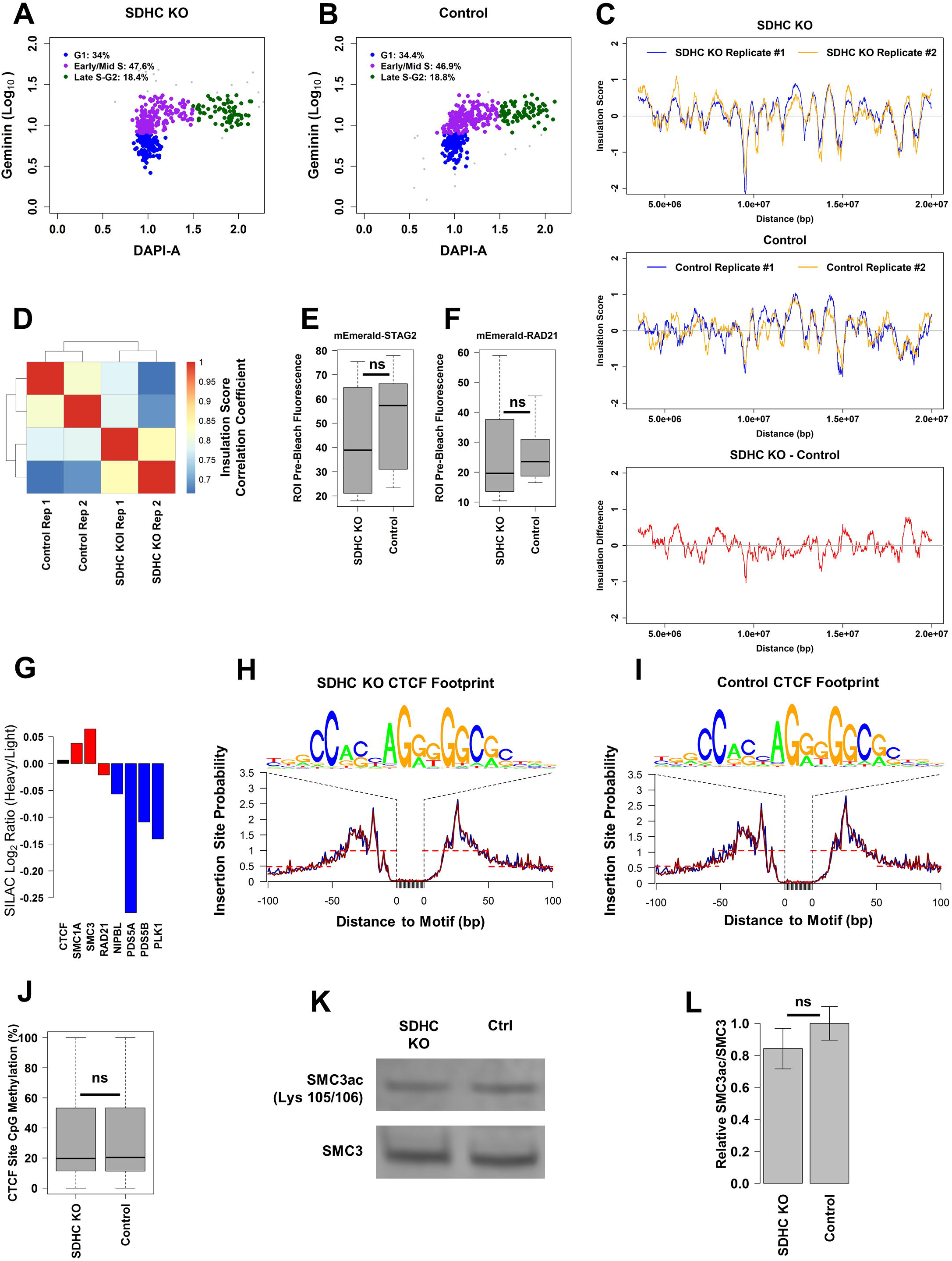
Supplemental analysis of cell cycle stages, factors affecting CTCF genomic occupancy, and cohesin acetylation, related to Figure 4. (A,B) Flow cytometry analysis of cell cycle stages in SDHC KO and control cells by DAPI-Geminin staining. (C) Insulation scores calculated for first 20 Mbp of chromosome 1 for biological replicate experiments for SDHC KO and control cells. (D) Heatmap showing genome-wide correlation for insulation between insulation scores calculated for biological replicate SDHC KO and control cells. (E,F) Analysis of initial cellular fluorescence for ROIs in FRAP experiments using mEmerald-STAG2 (E) and mEmerald-RAD21 (F). Statistical comparison of distributions is performed using a Wilcox rank-sum test. (G) Quantifications of change in expression of CTCF (black), cohesin subunits (red) and known cohesin regulators (blue) in SDHC KO relative to control cells by SILAC proteomics. (H,I) Genome-wide CTCF footprints calculated from ATAC-seq data for SDHC KO and control cells. (J) Quantification of CpG methylation at CTCF binding sites in SDHC KO and control cells (“ns” denotes lack of statistical significance in comparison of distributions by Wilcox rank-sum test). (K) Western blot analysis of SMC3ac Lys105/106 and total SMC3 in SDHC KO and control cell lines. (L) Quantitative analysis of SMC3ac Lys 105/106 relative to total SMC3. Statistical comparison by unpaired two-tailed t-test.

## STAR methods

### Immortalized mouse embryonic fibroblast cell lines

SDHC KO and hemizygous control immortalized mouse embryonic fibroblast cell lines used in this study have been described previously (Smestad et al., 2018a; Smestad et al., 2018b; Smestad and Maher, 2019). For routine cell culture applications, cells were grown in DMEM containing 10% FBS (Gibco cat# 10437), penicillin/streptomycin antibiotics (0.5 mg/mL), non-essential amino acids (100 µM each of glycine, alanine, asparagine, aspartic acid, glutamic acid, proline, and serine), sodium pyruvate (1 mM), and HEPES buffer (10 mM) at 21% O_2_ and 5% CO_2_.

### Metabolite extraction

Metabolite extraction was performed as described in a previous study (Liu et al., 2015). Briefly, SDHC KO and control iMEF cells were plated in triplicate into 6-well plates at a concentration of 5 million cells per well in DMEM containing 10% FBS (Gibco cat# 10437), penicillin/streptomycin antibiotics (0.5 mg/mL), non-essential amino acids (100 µM each of glycine, alanine, asparagine, aspartic acid, glutamic acid, proline, and serine), sodium pyruvate (1 mM), and HEPES buffer (10 mM). After 20 h, media was aspirated and the cells were washed with 0.9% NaCl. All residual saline solution was then aspirated and the plate was placed on dry ice. Pre-chilled 80% methanol/water (1 mL; HPLC-grade methanol: Sigma Aldrich cat# 34860-1L-R; HPLC-grade water: Sigma Aldrich cat # 34877-1L) was added to each well, and the plate was transferred to the −80°C freezer for 15 min for further inactivation of enzyme activities. Cells were then scraped into the methanol/water mixture on dry ice and transferred to 1.5 mL Eppendorf microcentrifuge tubes. Samples were subjected to centrifugation at 20,000×g for 10 min at 4°C to pellet cellular debris, and the clarified supernatant solutions were then transferred into new 1.5 mL Eppendorf microcentrifuge tubes prior to drying of samples on a Speed Vac at room temperature. Dried pellets were then stored at −80°C until use in downstream analysis. The dry pellets were reconstituted into 30 µL sample solvent (water:methanol:acetonitrile, 2:1:1, v/v) and 3 µL was further analyzed by liquid chromatography-mass spectrometry (LC-MS).

### Metabolite LC-MS

Ultimate 3000 UHPLC (Dionex) is coupled to Q Exactive Plus-Mass spectrometer (QE-MS, Thermo Scientific) for metabolite profiling. A hydrophilic interaction chromatography method (HILIC) employing an Xbridge amide column (100 × 2.1 mm i.d., 3.5 µm; Waters) is used for polar metabolite separation. Detailed LC method was described previously (Liu et al., 2014) except that mobile phase A was replaced with water containing 5 mM ammonium acetate (pH 6.8). The QE-MS is equipped with a HESI probe with related parameters set as below: heater temperature, 120 °C; sheath gas, 30; auxiliary gas, 10; sweep gas, 3; spray voltage, 3.0 kV for the positive mode and 2.5 kV for the negative mode; capillary temperature, 320 °C; S-lens, 55; A scan range (m/z) of 70 to 900 was used in positive mode from 1.31 to 12.5 min. For negative mode, a scan range of 70 to 900 was used from 1.31 to 6.6 min and then 100 to 1,000 min; resolution: 70000; automated gain control (AGC), 3 × 10^6^ ions. Customized mass calibration was performed before data acquisition.

### Metabolomics data analysis

LC-MS peak extraction and integration were performed using commercial available software Sieve 2.2 (Thermo Scientific). The peak area was used to represent the relative abundance of each metabolite in different samples. The missing values were handled as described in a previous study (Liu et al., 2014). Statistical testing for differential metabolite abundance was performed using two-sided t-test with FDR correction for multiple hypothesis testing. Chemical structural similarity analysis was performed with ChemRICH (Barupal and Fiehn, 2017).

### NAD^+^/NADH ratio quantification

SDHC KO and control cells were plated at a density of 3 × 10^5^ cells per well of a 6 well plate and allowed to recover overnight. The next day, cells were dissociated by trypsinization, quenched with media, pelleted by centrifugation for 5 min at 500×g, and washed with cold PBS. Measurement of NAD^+^/NADH ratio was performed using the BioVision NAD^+^/NADH quantification colorimetric kit (BioVision catalog# K337). Statistical testing was performed using a two-tailed t-test.

### Sirtuin activity measurements

Measurement of sirtuin activity in SDHC KO and control iMEF cell lines was according to a previously-published method (Xuan et al., 2017). pcDNA-mCherry-EGFP (K85TAG) and pCMVAcKRS-tRNAPyl plasmids used this experiment were the gifts of Peter Schultz (Scripps Research Institute). Briefly, SDHC KO and control iMEF cells were seeded at 30,000 cells/well in 12-well plates under the standard cell culture conditions specified above. The next day, pcDNA-mCherry-EGFP (K85TAG) and pCMVAcKRS-tRNAPyl plasmids were diluted into Opti-MEM, combined with diluted Lipofectamine 3000 reagent, and co-transfected into cells. One h later, acetyllysine solution was added to the wells to final concentration of 5 mM, and the plate was returned to the incubator overnight. The next day, mCherry and eGFP signals in individual cells were analyzed by analytical flow cytometry. Relative sirtuin activity was quantified by measuring the degree of eGFP fluorescence in the subset of dual mCherry and eGFP-positive cells for each cell line. Statistical testing for differences in sirtuin activity was performed using a Wilcox rank-sum test.

### SILAC proteomic quantification of acylation

Previously published SILAC proteomic datasets for SDHC KO and control cells (Smestad et al., 2018a; Smestad et al., 2018b) were searched for quantifiable site-specific acylation modifications. Modification SILAC H/L ratios were subsequently normalized according to protein abundance in the respective cell lines. For statistical testing, normalized SILAC H/L ratios for all quantified sites were grouped according to acylation type, and distributions of SILAC H/L ratios were compared by 1-way ANOVA for differences between categories.

### Western blotting

For whole cell extraction, SDHC KO and control cells were lysed in RIPA buffer (50 mM Tris-HCl, 5 mM EDTA, 150 mM NaCl, 0.1% SDS, 0.5% DOC, 1% NP-40) on ice for 15 min, with gentle pipetting and intermittent vortexing. Lysates were then centrifuged at 15,000×g for 5 min to pellet cellular debris, and protein concentrations of the recovered supernatants were quantified by BCA assay.

For analysis of subcellular fractions, SDHC KO and control cells harvested by trypsinization, pelleted via centrifugation at 600×g for 5 min at 4°C, washed 2× with ice cold PBS, and washed once with ice cold TBS. Following the last wash, cells were lysed in 0.3 mL hypotonic lysis buffer (10 mM Hepes-NaOH pH 7.9, 10 mM KCl, 1.5 mM MgCl_2_, 0.5 mM DTT, 0.1% (v/v) NP-40, and 1× Halt protease inhibitor) for 10 min with gentle agitation at 4°C. Nuclei were pelleted by centrifugation at 720×g for 5 min, and supernatant containing the cytosolic and membrane fraction was transferred to a new tube. Pelleted nuclei were resuspended in 200 µL of TBS containing 0.1% SDS and sonicated briefly to shear the genomic DNA and to solubilize nuclear proteins. Protein concentrations from the respective cytosolic and nuclear subcellular fractions were determined using the BCA assay.

Total protein (60 µg) was then combined with 4× LDS buffer including reducing agent, and heated to 90°C for 5 min to denature the proteins. Samples were then loaded onto a 10% bis-tris gel (NuPAGE, cat# NP0301BOX) and subjected to electrophoresis at 150 V for 35 min in 1× MES-SDS running buffer. Blot transfer onto a PVDF membrane was conducted at 30V for 90 min at 4°C in NuPAGE transfer buffer containing 20% methanol. Following transfer, proteins were visualized on the membrane using Ponceau S stain. Membranes were then blocked for 1 h at room temperature in blocking buffer comprised of TBST containing 3% non-fat dried milk. Membranes were then washed 3× 5 min with 1× TBST. Primary antibody solutions were prepared in antibody dilution buffer containing 2.5 mL 4% BSA, 250 µL 0.5% NaAzide, 7.5 mL TBST. Antibodies used in this analysis include anti-SDHC rabbit polyclonal IgG antibody (Santa Cruz Biotechnology, cat# sc-67256 (M-169), 1:500 dilution), anti-actin rabbit polyclonal IgG antibody (Sigma, cat# A2066, 1:500 dilution), anti-SIRT1 rabbit monoclonal IgG antibody (Cell Signaling Technology cat# 9475, 1:1000 dilution), anti-Phospho SIRT1 (Ser47) rabbit monoclonal IgG antibody (Cell Signaling Technology cat# 2314, 1:2000 dilution), anti-SIRT2 rabbit monoclonal IgG antibody (Cell Signaling Technology cat# 12650, 1:1000 dilution), anti-SIRT3 rabbit monoclonal IgG antibody (Cell Signaling Technology cat# 5490, 1:1000 dilution), anti-SIRT5 rabbit monoclonal IgG antibody (Cell Signaling Technology cat# 8782, 1:1000 dilution), anti-SIRT6 rabbit monoclonal IgG antibody (Cell Signaling Technology cat# 12486, 1:1000 dilution), anti-SIRT7 rabbit monoclonal IgG antibody (Cell Signaling Technology cat# 5360, 1:1000 dilution), anti-histone H3K27ac rabbit polyclonal IgG antibody (Novus Biologicals cat# NBP2-54615, 1:1000 dilution), anti-histone H4ac (H4K5ac, H4K8ac, H4K12ac, and H4K16ac) rabbit monoclonal IgG antibody (Abcam cat# ab177790, 1:20000 dilution), pan anti-propionyl rabbit polyclonal antibody (PTM Biolabs cat# PTM-201), pan anti-butyryl rabbit polyclonal antibody (PTM Biolabs cat# PTM-301), pan anti-malonyl rabbit polyclonal antibody (PTM Biolabs cat# PTM-901), pan anti-succinyl rabbit polyclonal antibody (PTM Biolabs cat# PTM-401), anti-histone H4 rabbit polyclonal IgG antibody (Abcam cat# ab7311, 1:1000 dilution), and anti-histone H3 rabbit polyclonal IgG antibody (Abcam cat# ab1791, 1:1000 dilution). Following dilution of antibodies into antibody dilution buffer, solutions were added to the membranes and incubated at 4°C overnight with gentle shaking. The next day, solutions of primary antibodies were removed and membranes were washed 3× 5 minutes with 1× TBST at room temperature. Membranes were then incubated with HRP-conjugated anti-rabbit IgG secondary antibodies, diluted 1:10,000 into blocking buffer, for 1 h at room temperature. Membranes were then washed 3× 5 min with TBST to remove excess antibody, and exposed to ECL2 Western blot substrate for 5 min at room temperature prior to fluorescence image acquisition using a Typhoon fluorimeter.

### Transmission electron microscopy

SDHC KO and control cells were fixed in EM fixative (4 % paraformaldehyde with 1% glutaraldehyde in phosphate buffered saline, pH 7.2), and placed into 2% low melting agar. Cells were then stained with 1% osmium tetroxide and 2% uranyl acetate, dehydrated through an ethanol series, and embedded into Spurr resin. Following a 24-h polymerization at 60°C, 0.1 µm ultra-thin sections were post-stained with lead citrate. Micrographs were acquired using a JEOL1400 Transmission Electron Microscope (Peabody, MA) operating at 80kV.

### CUT&RUN

CUT&RUN sequencing was performed according to a published protocol (Skene et al., 2018). pA-MN fusion protein was gifted by Steven Henikoff (Fred Hutchinson Cancer Research Center, HHMI). Antibodies used for this experiment include anti-CTCF rabbit polyclonal antibody (Abcam cat# ab70303), anti-H3K4me3 rabbit polyclonal antibody (Abcam cat# ab8580), anti-H3K27ac rabbit polyclonal antibody (Active Motif cat# 39134), anti-H3K27me2 rabbit polyclonal antibody (Abcam cat# 24684), pan anti-acetyllysine rabbit polyclonal antibody (PTM Biolabs cat# PTM-105), pan anti-propionlyllysine rabbit polyclonal antibody (PTM Biolabs cat# PTM-201), and pan anti-butyryllysine rabbit polyclonal antibody (PTM Biolabs cat# PTM-301). Following recovery of eluted chromatin fragments CUT&RUN, library preparation was completed using the ThruPLEX DNA-seq 48S kit (Rubicon Genomics cat# R400427) and purified using AMPure XP beads (Beckman Coulter cat# A63880). 40 bar-coded libraries were multiplexed into a single lane on an Illumina HiSeq 4000 instrument and paired-end index sequencing performed to 150-bp read length. FASTQ reads were then aligned to the GRCm38/mm10 mouse reference genome using Bowtie2 on the Mayo Clinic Research Computing Services high performance Beowulf-style Linux-based cluster. Chromatin epigenomic state discovery was performed in ChromHMM (Ernst and Kellis, 2017). Data were deposited at GEO NCBI and are publically available under identifier GSE129956.

### ATAC-seq

OMNI ATAC-seq was performed for 50,000 cells following the previously-published protocol (Corces et al., 2017). Amplified libraries were purified, and the size distribution of library DNA was determined by Fragment Analyzer (Advanced Analytical Technologies, IA) using a High Sensitivity NGS Fragment Analysis Kit (cat# DNF-486). Enrichment of accessible regions was determined by quantification of fold difference between positive and negative genomic loci using real-time PCR. Primer pairs for positive loci include AT-mDusp6-F: 5’-GGCTTATCCGGAGCGGAAAT and AT-mDusp6-R: 5’-GGCTGGAACAGGTTGTGTTG, and AT-mCh8-F: 5’-ACAAACATGCAGCAAGCCAC and AT-mCh8-R: 5’-ACTCACTGGCCAATCAAGGC-3’. Primers for negative locus amplification include AT-mVmn2r17-F: 5’-TCCCCTTTACTGTTTTCCTCTAC and AT-mVmn2r17-R: 5’-GGATTGATGAGGAAACAGCCTC. Four bar-coded libraries were multiplexed into a single lane on an Illumina HiSeq 4000 instrument and paired-end index sequencing performed to 50-bp read length. FASTQ reads were then aligned to the GRCm38/mm10 mouse reference genome using Bowtie2 on the Mayo Clinic Research Computing Services high performance Beowulf-style Linux-based cluster. ATAC-seq peak calling was performed in SeqMonk (https://www.bioinformatics.babraham.ac.uk/projects/seqmonk/) using the MACS peak calling algorithm (p-value cutoff: 1E-5), performing separate analyses for SDHC KO and control cell lines. Nucleosome positioning analysis of ATAC-seq data was performed using the NucleoATAC pipeline (Schep et al., 2015) for regions mapping to chromatin states determined by CUT&RUN/ChIP-seq and ChromHMM (Ernst and Kellis, 2017). Inter-dyad distances were calculated from the positions of called nucleosomes. Statistical testing for differences between groups was performed using a Wilcox rank-sum test. Inferred CTCF occupancy at CTCF binding sites was calculated using the ATACseqQC R package (Ou et al., 2018). Data were deposited at GEO NCBI and are publically available under identifier GSE129956.

### DNA template preparation and cloning

Plasmid pET-15b-derived vectors for expression of *Homo sapiens* histones H2A (pHISPP/H2A), H2B (pHISPP/H2B), H3.1 (pHISPP/H3.1), and H4 (pHISPP/H4) were the gifts of Georges Mer. The sequence identifiers/internal plasmid codes for these constructs are NP_003504.2, NP_066402.2, NP_003520.1, and NP_001029249.1, respectively). Plasmid pJ2497 was used to create a transcription template for a T7 RNA polymerase transcription elongation assay in the presence or absence of reconstituted nucleosomes. This plasmid was propagated by standard methods in XL-10 Gold *E. coli* cells (Agilent) after heat shock transformation. Single colonies were grown on LB agar plates containing carbenicillin (50 µg/mL), transferred to 5 mL LB medium containing carbenicillin (50 µg/mL), and grown at 37°C for 16-20 h with continuous shaking at 225 rpm. The resulting culture was then used to seed 1 L of LB medium containing carbenicillin (50 µg/mL) and grown at 37°C for 16-20 h with continuous shaking at 225 rpm. Plasmid DNA was isolated by DNA maxi preparation kit (Qiagen # 12163) using 125 mL culture per tip-500 column. To generate the 624-bp transcription template for nucleosome reconstitution and T7 RNA polymerase transcription, plasmid pJ2497 DNA was subjected to restriction endonuclease digestion with *Pci*I (NEB # R0655) using 140 U enzyme per mg plasmid in NEB buffer 3.1 at 37°C for 10-14 h. The resulting digest was diluted to 300 ng/µL, subjected to electrophoresis through a 0.8% agarose gel for 2.5 h at ∼6 V/cm, and the DNA band corresponding to the 624-bp transcription template was purified from the gel using a QIAquick Gel Extraction Kit (Qiagen #28704). The gel purified 624-bp transcription template DNA was further purified by ethanol precipitation involving addition of 0.1 volume of 3 M NaOAc, pH 5.2 and 3 volumes of 100% ice cold ethanol with cooling at −80°C for 1 h. The ethanol precipitated DNA was then subjected to centrifugation at 14,000 x g for 15 min at 4°C and the supernatant was discarded. Pelleted DNA was then washed with 750 μL 70% EtOH and subjected to centrifugation as before. The dried DNA was typically resuspended in 50 μ L water. The concentration of purified 624-bp transcription template was determined by ultraviolet absorbance measured at 260 nm.

### Preparation of histone octamers

Histone expression vectors were electroporated into BL21(DE3) cells and plated on LB agar plates containing carbenicillin (50 µg/mL). Transformed colonies were used to inoculate 1 L LB medium containing carbenicillin (50ug/mL). The resulting cultures were grown at 37°C with shaking at 225 rpm. When cultures reached an OD_600_ of 0.6, IPTG was added to a final concentration of 0.5 mM and growth continued for an additional 3 h at the same conditions. Cultures were subjected to centrifugation at 6000 × g for 10 min in centrifuge Beckman Coulter Rotor JLA 8.1000 and LB media was removed.

Pelleted cells were resuspended in 10 mL ice-cold resuspension buffer (50 mM potassium phosphate pH 7.5, 300 mM NaCl, 1 mM phenylmethylsulfonyl fluoride, PMSF) and lyzed by passage three times through a French press (AVESTIN Emulsi Flex-05) at 0.5 kPa. The lysate was subjected to centrifugation at 20,000 × g for 45 min at 4°C in (Beckman Coulter Rotor JA-25.50). The supernatant was discarded and the insoluble pellet, containing the histone proteins, was solubilized by the addition of 10 mL ice-cold binding buffer (50 mM potassium phosphate pH 7.5, 8 M urea, 300 mM NaCl, 5 mM imidazole, 1 mM PMFS). To aid in solubilization, histone proteins were then subjected to sonication at 30% power, three times at intervals of 30 s on ice (Branson sonifier) allowing 30-s incubation on ice between sonications to avoid overheating. The sonicated solution was then clarified by centrifugation at 20,000 g for 45 min at 4°C (Beckman Coulter rotor JLA-16.250). The resulting supernatant was then allowed to pass by gravity flow over a column containing 4 mL packed Ni-NTA beads (Qiagen #30230) previously equilibrated in binding buffer, and the resulting flow-through was discarded. Columns were washed with 300 mL wash buffer (50 mM potassium phosphate pH 7.5, 6 M urea, 300 mM NaCl, 20 mM imidazole, 1 mM PMSF) using a Longer Pump (BT100-IL) at a flow rate of 3.5 mL/min. Histone proteins were eluted from the column with 30 mL elution buffer (50 mM potassium phosphate pH 7.5, 6 M urea, 300 mM NaCl, 250 mM imidazole, 1 mM PMFS).

Concentrations of eluted histones were estimated by their absorbance at 280 nm using respective molecular weight and molar extinction coefficient values calculated using the Proto Param Tool (Wilkins et al., 1999). Concentration and quality of purified histones were validated empirically by electrophoresis through denaturing 10% Bis-Tris SDS polyacrylamide gels (NuPage #NPO363BOX) at 125 V for 2-3 h. Purified histone proteins in elution buffer were aliquoted, flash frozen on dry ice, and stored at −80°C.

Equimolar amounts of histones H2A, H2B, H3.1, H4 or (H4 K79 variants) were combined in varying volumes of elution buffer depending on their concentration. Dialysis steps were then performed at 4°C using 7 kDa MWCO SnakeSkin Dialysis tubing (Thermo SCIENTIFIC Lot#RF235431). Dialysis began against 2 L octamer refolding buffer 1 (10 mM Tris HCl pH 7.5, 4 M Urea, 2 M NaCl, 1 mM EDTA, 200 μM EDTA, 5 mM 2-mercaptoethanol) for 10-14 h. The concentration of urea in the refolding buffer was then decreased stepwise by 1 M every 6 h in a series of subsequent dialysis steps until dialysis against buffer containing 1 M urea had been completed. Histones were then dialyzed against octamer refolding buffer 2 (10 mM Tris HCl pH 7.5, 2 M NaCl, 1 mM EDTA, 200 μ M EDTA, 5 mM 2-Mercaptoethanol, for 10-14 h). Cleavage of His_6_ affinity tags from refolded histone octamers was accomplished by treatment with PreScission Protease (Sigma, SAE0045) for 10-14 h at 4°C. Histones were then dialyzed against fresh octamer refolding buffer 2 for 10-14 h at 4°C. Affinity tag cleavage was confirmed by SDS polyacrylamide gel electrophoresis with an extended run time of 4h. Refolded histone octamers were concentrated to a final volume of 2 mL using a 10 kDa MWCO concentrator (Vivaspin 20 # VS2002) subjected to centrifugation at 3800 rpm (Jouan T 1354 table top centrifuge).

Histone octamers were purified from other multimeric histone species by size exclusion chromatography using a GE ÄKTA FPLC instrument as described (Dyer et al., 2004). Fractions containing histone octamers were combined and concentrated as above to 1-2 mL. The concentration of histone octamer was estimated by absorbance at 280 nm using molar extinction coefficients calculated using the ProtParam tool (1). Octamers were stored at −20°C in 10 mM Tris-HCl pH 7.5, 50% glycerol, 2 M NaCl, 2 mM dithiothreitol.

### Nucleosome core particle (NCP) reconstitution

NCPs and a DNA-only control lacking histone octamer were reconstituted in parallel using stepwise dialysis of the purified 624-bp transcription template DNA and histone octamer at a molar ratio of 1:1.2, respectively. This was accomplished by dialysis of a solution consisting of 6 µg transcription template DNA, 2 µg histone octamer (added last, with mixing) in a final L solution containing 10 mM Tris-HCl pH 7.5, 2 M NaCl, 250 μ 2-Mercaptoethanol. The DNA-only solution was made in parallel, excluding histone octamer. Equilibration and dialysis steps were performed at 4°C on an orbital shaker (Fisher Scientific Clinical Rotator cat 14-251-200) at 60 rpm using Slide-A-Lyzer MINI dialysis 3.5kD MWCO units (Thermo-Fisher) against dialysis buffers containing 10 mM Tris-HCl pH 7.5, 2 M NaCl, 250 μ M EDTA, 5 mM 2-mercaptoethanol for 30 min. Reconstitutions were then dialyzed at 4°C against dialysis buffers containing 1.2 M NaCl (4-6 h), 0.85 M NaCl (4-6 h), 0.65 M NaCl (10-14 h), 0.5 M NaCl (4-6 h) and 2.5 mM NaCl (4-6 h). Solutions were then subjected to centrifugation at 14,000 × g for 15 min to pellet any precipitated material. The final concentration of transcription templates was estimated by the solution volume after dialysis assuming no loss of DNA.

### *In vitro* chemical acylation of reconstituted NCPs

Reconstituted nucleosomes were dialyzed into HEPES buffer to remove Tris-HCl. Acyl-CoA stocks were prepared by dissolving commercial powders (Sigma) in water at a final concentration of 20 mM. Stocks were generally prepared fresh. Stocks were added 1:1 to nucleosomes to 10 mM final concentration and allowed to react for 3 h at room temperature unless otherwise indicated. Treated reconstituted nucleosomal templates were then analyzed without further purification.

### Electrophoretic mobility shift assay of nucleosome reconstitution

A native 1.2% agarose electrophoretic mobility shift assay was used to estimate the extent of NCP reconstitution on the 624-bp DNA transcription template derive from plasmid pJ2497. For this assay, the reconstituted transcription template (∼600 ng DNA) was first subjected to digestion with *Hin*dIII-HF (NEB #R3104; 30 U) in CutSmart buffer (NEB) at 37°C for 1 h to release 375-bp and 249-bp fragments, the latter of which carries the NCP. Samples were subjected to electrophoresis through a 1.2% native agarose gel for 2.5 h at ∼6 V/cm. The gel was then stained for 40 min in a solution of 40 mM Tris-HCl, 20 mM acetic acid, 1mM EDTA containing ethidium bromide (0.75 µg/mL), and de-stained for 20 min in the same solution without ethidium bromide. Gel fluorescence was then detected on a Typhoon FLA 7000 imager using a 610-nm band-pass filter. The fraction of template fragment reconstituted with NCP was estimated using NIH ImageJ software.

### T7 RNA polymerase transcription assay through reconstituted nucleosome

*In vitro* transcription reactions to assay nucleosome stability were performed by mapping radiolabeled RNA transcription products of T7 RNA polymerase elongation into a reconstituted nucleosome. Initial *in vitro* transcription labeling reactions contained reconstituted transcription templates (125 ng DNA) in T7 RNA polymerase buffer including CTP and GTP (50 µM each), and α-[^32^P] ATP (800 Ci/mmol; 1μM), dithiothreitol (5 μΜ), 1.5 μL RNasin (N2111; 1 U/μL), and 2.5 U/μL T7 RNA polymerase (NEB M0251L), in a final volume of 50 μL. Transcription was performed at room temperature for 10 min to radiolabel synthesized RNAs in the T-free template region. Samples (20 µL) from the zero time point were collected from ice-cold reactions and added to an equal volume of deionized formamide and boiled at 90°C for 3 min. To initiate transcription elongation beyond the T-free template region, 25 mM NTPs were added to 2 mM final concentration and transcription was allowed to proceed for 1 min at 4°C. Samples (20 µL) were collected and terminated as above. A reference ladder of radiolabeled RNA transcripts was transcribed in the same manner using a mixture of templates individually digested by restriction enzymes PciI (NEB RO655,) NotI (NEB RO189), RsaI (NEB RO167), PmlI (NEB RO532), or XbaI (NEB RO145). Transcription reaction products were then subjected to electrophoresis through polyacrylamide gels (29:1 acrylamine:bisacrylamide) containing 7.5 M urea at 12 W for 1 h. A 20% volume of 2.5 M sodium acetate was then added the lower electrophoresis reservoir with electrophoresis at 12 watts for an additional 45 min to enhance band separation. The gel was then dried and analyzed by storage phosphor imaging technology corresponding to polymerase pausing or termination near the NCP were mapped by regression fitting of RNA markers.

### Quantitation of nucleosome reconstitution and transcription pausing/termination

The fraction (*N*) of DNA template reconstitution into nucleosomes was calculated (after background correction) by comparing the unshifted 601 (*N*) and non-601 (*T*) fragment intensities after nucleosome reconstitution to the corresponding values (N’, T’) in the absence of histone octamer:

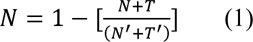

The fraction (*B*) of transcription pause and termination was calculated by densitometry measurement (after background correction) of the full-length transcript band (*F*) and the cluster of transcript pause bands (*P*):

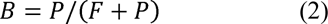

The overall Nucleosome Obstacle Activity Per Reconstituted Template (NOAPRT) parameter was then calculated:

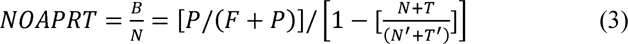

### Nucleosome arrays were assembled for optical tweezers

experiments as described previously (McCauley et al., 2019). The DNA template was constructed around the 147-bp octamer positioning 601 sequence joined to a 60-bp non-specific linking sequence (Lowary and Widom, 1998). A linear 12× array of these 207-bp repeats was attached to linker DNA for optical tweezers experiments by insertion into pUC19 plasmid DNA. Endonuclease digestion (*Bsa*I) linearized this construct and left a 4-bp overhang on both termini, filled in with DNA polymerase I (*Klenow*) and digoxygenin-labeled dTTP and biotinylated dATP (as well as dGTP and dCTP). The resulting construct consisted of a 12× array of 207-bp nucleosome sites, flanked by ∼1350-bp non-positioning DNA linkers, and ended with a single biotin and digoxygenin each on opposing termini. DNA human octamers were assembled onto this construct by salt titration/dialysis (Muthurajan et al., 2011; Rogge et al., 2013). Essentially, a high concentration of DNA construct described above (∼0.40 μg/μL) was incubated in high salt (10 mM Tris–HCl, pH 7.5, 1 mM EDTA and 2 M Na^+^). Human octamers (gift of the Lugar lab at the University of Colorado) were introduced into this solution in excess to ensure full array assembly. This solution was transferred into a 50 μL volume dialysis button and dialyzed (Hampton) against a series of decreasing salt solutions over ∼ 30 h, facilitating nucleosome assembly at each positioning site. Arrays were stored in the final buffer (10 mM Tris–HCl, pH7.5, 1 mM EDTA and 2.5 mM Na^+^), and remained stable over several weeks at 4 °C. Reconstituted nucleosome arrays were assessed via atomic force microscopy (AFM) imaging and optical tweezers force microscopy (OT). OT experiments verify that 11-12 nucleosomes were disrupted by force and are described below. AFM experiments imaged arrays in liquid solutions to verify successful nm, and clusters of 10-12 nucleosomes were seen in stable arrays.

### Single molecule optical tweezers experiments

The dual beam, single trap optical tweezers is well-known and widely-used (Chaurasiya et al., 2010; McCauley and Williams, 2009). Prior to experiment, reconstituted nucleosome arrays were diluted 10,000:1 into a solution of 10 mM HEPES, pH7.5 and 100 mM Na^+^ (a final array concentration of 0.030 nM). An experiment began with the ‘catch’ of a single nucleosome array between a 2.1-μm diameter anti-digoxygenin coated bead (*Spherotech*) and a 3.4-μm diameter streptavidin-coated bead (*Bangs Labs*). The smaller bead was pulled onto a micropipette tip (*WPI*) within a custom-built flow cell. The larger bead was held at the focus an optical trap of finite stiffness. Movement of the cell altered the tension in the array, which is measured at each step. Experiments typically began at low extension and increased stepwise at a nearly fixed loading rate of 10 pN/s, followed by a stepwise decrease in extension.

Arrays were diluted and experiments were performed at room temperature (∼22 °C). Control experiments stretched arrays alone and though some free octamers were present in solution, there was no evidence of non-specific binding to the long DNA handles. For the control, arrays with *N* > 10 observed disruptions were selected. Diluted array samples were also separately incubated with 2.0 mM acetyl-CoA, propionyl-CoA, butyryl-CoA, malonyl-CoA and succinyl-CoA at 30 °C for 3 h. Solutions of incubated arrays were then introduced directly into the optical tweezers experiment (22 °C) and tethered for cycles of extension and release. Arrays with *N* > 7 nucleosome disruptions were retained. A further control was performed where diluted disruption by exposure to temperature in a dilute solution (Hazan et al., 2015).

### Polymer elasticity modeling

The elastic response of double-stranded DNA has been well-studied under a variety of conditions. With the application of an external tension, the end-to-end distance of a flexible worm-like chain will respond as (Bouchiat et al., 1999; Chaurasiya et al., 2010; McCauley et al., 2013; McCauley and Williams, 2009),

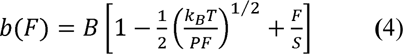

Adjustable parameters include the contour length *B,* the elastic modulus *S* and the persistence length *P*, a key measure of chain flexibility (Chaurasiya et al., 2010; McCauley et al., 2013; Murugesapillai et al., 2014). There is known variation of these parameters upon construct length (Seol et al., 2007) and for these constructs (without nucleosomes) fitted values are *B* = 0.340 ± 0.001 nm/bp (or 1782 ± 5 nm), *S* = 800 ± 100 pN and *P* = 40 ± 2 nm. Thus, the contour length for a fully-reconstituted construct would reasonably be (1782 – 78×*N*) nm, where *N* is the number of nucleosomes remaining in the array (as defined in the main text).

DNA constructs (nucleosome free) were incubated with both acetyl- and succinyl-CoA. Fits to the polymer elasticity model described above quantify ligand binding that are known to alter either the length of the double strand or the local flexibility (Chaurasiya et al., 2010; McCauley et al., 2019; McCauley et al., 2013). Non-linear fits are written into custom LabWindows CVI code which incorporated a ML Numerical Recipes algorithm (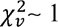 for these fits) (Press et al., 2002).

### Kinetics of nucleosome unwinding

As DNA wrapped around a single nucleosome is subjected to increasing tension, there is an increasing probability that the nucleosome will be disrupted. Arrays of multiple nucleosomes will show disruptions with increasing force as the probability of observing any individual disruption increases with the decreasing number left in the array, *N* (McCauley et al., 2019),

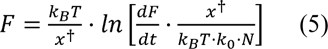

The loading rate, *dF*/*dt,* is constant over all the experiments here (∼10 pN/s). This kinetic information allows us to characterize the transition barrier for strong site ripping. Specifically, we quantify the DNA extension to the transition state (*x^†^*) and the natural (zero force) rate of unwinding (*k_o_*) are fitting parameters. However, for nucleosome arrays, the energy landscape is complicated by many likely sub-transition states, so these two parameters were fitted only for the control set (which found *x^†^* = 0.47 ± 0.01 nm, or 1.38 ± 0.03 base pairs), then held fixed for the remaining experiments (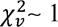 for all fits).

### Easy Hi-C

Easy Hi-C was performed as previously reported using 1×10^6^ cells as input (Lu et al., 2018). Library preparation using 50 ng recovered DNA was completed using the ThruPLEX DNA-seq 48S kit (Rubicon Genomics cat# R400427) and purified using AMPure XP beads (Beckman Coulter cat# A63880). Four bar-coded libraries were multiplexed into a single lane on an Illumina HiSeq 4000 instrument and paired end index sequencing performed to 150-bp read length. FASTQ reads were then analyzed on the Mayo Clinic Research Computing Services high performance Linux cluster using the Juicer Hi-C analysis pipeline (Durand et al., 2016). All downstream analyses employed Knight-Ruiz matrix balancing for contact map normalization (Knight and Ruiz, 2013). Analysis of eHi-C dataset variability between biological replicates was performed using 1-Mb resolution contact matrices in HiCRep (Yang et al., 2017). Eigenvectors used to delineate chromatin compartments at 250-Kb resolution were calculated in Juicer tools (Durand et al., 2016). Identities of A and B compartments were determined by assessing correlation between compartment eigenvectors and binned ATAC-seq signal. TAD calling using insulation square analysis and calculation of TAD boundary strength was performed as previously described on a 10-Kbp resolution contact matrix using a sliding 500 Kbp × 500 Kbp (50 bin × 50 bin) (Crane et al., 2015). TADs retained for analysis have mean boundary strength > 0.5 across all datasets. TADs with differential boundary strength were those identified to have absolute log_2_(fold-change) > 1. Data were deposited at GEO NCBI and are publically available under identifier GSE129956.

### Cell cycle analysis

SDHC KO and control cells were dissociated by trypsinization, resuspended in media, diluted to 1×10^6^ cells/mL, subjected to centrifugation at 500×g for 5 min, resuspended in PBS containing 2% formaldehyde, and rocked for 20 min at room temperature to fix. The formaldehyde reaction was quenched with the addition of glycine to 127 mM final concentration and incubating on ice for 5 min. Cells were subjected to centrifugation at 500×g for 5 min, washed 2× with PBS, resuspended in permeabilization buffer (10 mM Tris-HCl (pH 8.0), 10 mM NaCl, 0.2% IGEPAL CA-630), and incubated on ice for 30 min to permeabilize. Cells were then centrifuged at 500×g for 5 min, resuspended in blocking buffer (PBS containing 5% FBS and 0.1% Tween-20), and incubated at room temperature for 1 h. Cells were then subjected to centrifugation at 500×g for 5 min, resuspended in blocking buffer containing rabbit anti-Geminin polyclonal antibody (Abcam cat# ab175799) at 1:400 dilution, and incubated at 4°C with gentle agitation for 1 h. Cells were then subjected to centrifugation at 500×g for 5 min, washed with blocking buffer, and resuspended in blocking buffer containing goat anti-rabbit IgG (H+L) Alexa Fluor Plus 488-conjugated antibody (Invitrogen cat # A32731) at 1:200 dilution, and incubated at 4°C with gentle agitation for 30 min. Cells were then subjected to centrifugation at 500×g for 5 min, washed with blocking buffer, resuspended in PBS containing 0.1% TX-100 and 10 µg/mL DAPI, and incubated on ice for 30 min. Cells were then subjected to centrifugation at 500×g for 5 min and resuspended in PBS prior to flow cytometry analysis of DAPI and Geminin.

### Fluorescence recovery after photobleaching

mEmerald-STAG2-C-18 and mEmerald-Rad21-C-18 were gifts from Michael Davidson (Addgene plasmid # 54259 and 54247). Plasmids were propagated in DH5alpha with Kanamycin selection and isolated using Qiagen Plasmid Maxi kit (Qiagen cat # 12163). SDHC KO and control iMEF cells were seeded at 100,000 cells per 2 mL media into 30 mm plates under the standard cell culture conditions specified above. The next day, mEmerald-STAG2-C-18 or mEmerald-Rad21-C-18 plasmids were diluted into Opti-MEM, combined with diluted Lipofectamine 2000 reagent (Invitrogen cat # 11668030), and transfected into cells. The media was changed 4 h later and the plate was returned to the incubator overnight. The next day, cells were analyzed by confocal microscope using a 488 nm laser and detection wavelengths of 493-598 nm in a fluorescence recovery after photo bleaching (FRAP) experiment utilizing a circular bleach area with 1 µM radius. Intensity measurements for regions of interest (ROIs) including intracellular bleached area, intracellular non-bleached area, and extracellular background area were captured at 1 s time intervals for 120 s. Extracellular background was subtracted from bleached and non-bleached intracellular ROIs, and then intensity ratio of intracellular bleached ROI to intracellular non-bleached ROI was calculated as a function of time to yield a normalized FRAP curve.

### Proteomic analysis of cohesin subunit acyl PTMs

SILAC heavy and light DMEM media were prepared using the SILAC Protein Quantitation Kit (LysC) DMEM (Thermo Scientific cat # A33969) and L-Arginine, ^13^C_6_ for SILAC (Thermo Scientific cat # 88210), with the addition of glutamine and pyruvate. Control cells were grown for 10 doublings in SILAC heavy media containing ^13^C_6_ L-Lysine and ^13^C_6_ L-Arginine, and SDHC KO cells were grown in SILAC light media containing ^12^C_6_ L-Lysine and ^12^C_6_ L-Arginine. 20 million cells from each line were trypsinized and washed 2× with cold PBS. Cells were lysed with 1500 µL cold lysis buffer containing 50 mM Tris-HCl, pH 7.4, 150 mM NaCl, 1% TX-100, and 1X protease inhibitor cocktail. Lysates were subjected to centrifugation at 15,000 rpm for 5 min at 4 °C and the supernatant transferred to a new tube. Protein concentration was estimated by BCA assay and lysates were diluted to a concentration of 1.5 mg/mL in 3 mL total volume. MgCl_2_ was added to 0.4 mM concentration and 10 µL DNase I (5 mg/mL) was added. Tubes were incubated at room temperature for 10 min to degrade genomic DNA. EDTA was added to final concentration of 5 mM and tubes were placed on ice to stop nuclease reaction. Samples were split into three technical replicates, combined with 25 µL of pre-washed protein A/G magnetic beads (Pierce cat # 88803) and 1 µg of non-specific rabbit IgG antibody, and rotated at 4 °C for 1 h. Samples were placed on magnet to separate beads and supernatant containing pre-cleared lysate transferred to a new tube. Rabbit anti-SMC3 antibody (8 µg; Abcam cat # ab9263) was added and tubes were again rotated at 4 °C for 2 h. 25 µL of washed protein A/G magnetic beads was added and tubes were rotated at 4 °C overnight to capture antibody-bound cohesin. Tubes were placed on a magnet to phase separate, and the beads were washed 2× with lysis buffer. Beads were resuspended in 50 µL of 1× LDS loading buffer containing sample reducing agent and heated to 95 °C for 5 min to elute bound proteins. 20 µL eluted sample was then subjected to electrophoresis on a 10% SDS-PAGE gel for 1 h at 130 V. The gel was then stained in Coomassie solution and de-stained in methanol-acetic acid solution. Bands corresponding to the expected size of cohesin subunits were excised using a clean razor blade. The bands with light and heavy labeled proteins were excised and pooled roughly equally for combined in-gel digestion by trypsin (Promega, Madison, WI) based on a previously described protocol (Li et al., 2019). Extracted peptides were desalted and subject to LC-MS/MS analysis. Peptides were separated on an in-house packed microcapillary C18 column and eluted with HPLC gradient of 1% to 30% HPLC buffer A (0.1% formic acid in water) in HPLC buffer B (0.1% formic acid in acetonitrile) from 5 min to 45 min. Peptide ions were acquired by Orbitrap Lumos Hydrid mass spectrometer (Thermofisher, Waltham, MA). Precursor ions were analyzed in Orbitrap with a resolution of 50,000 at 200 m/z. Dynamic exclusion was enabled. Peptide ions were fragmented through high-energy collision dissociation (HCD) and analyzed by ion trap.

Three biological replicate analysis of SILAC-labeled Cohesin complex proteins were searched with Maxquant software against the protein database containing Cohesin proteins and known contaminants. Common lysine acylations, including acetylation, succinylation, malonylation, propionylation, butyrylation, crotonylation, glutarylation, were specified as variable modifications in addition to oxidation on methionine and acetylation on protein N-terminus. Carbamidomethylation on Cys was specified as the fixed modification. Up to four missed cleavage in trypsin digestion were allowed. Precursor ion mass tolerance was set to 4.5 ppm and fragment ion mass tolerance was set to 0.6 Da. Protein quantification considered unmodified peptides and only peptides with oxidation on Met or acetylation on protein N-terminus. A false discovery rate of 1% was applied for protein, peptide and modification site identifications. Identification of modified peptides were also subject to a cutoff Andromeda score of 40. Quantification of PTM sites included the normalization of SILAC ratios of PTM sites with the corresponding protein SILAC ratios.

For PTM sites that could not be quantified by SILAC (missing heavy or light isotope-labeled peptide ions), we provided an estimation of the SILAC ratios based on a previous described strategy for data analysis with missing values (Ogden, 2010). Based on this strategy, we first estimated the MS detection limit (D) when the target peptide was eluting at its chromatographic peak. The MS detection limit (D) was calculated as the average intensity of the lowest one-third of the MS signals in the vicinity (+/- 100 m/z) of the detectable SILAC peptide ion. Then, we assigned an arbitrary value of 0.707xD as the missing intensity of the other SILAC peptide ion. As the MS intensity of the detectable SILAC peptide ion was known, this would allow us to estimate the upper or lower limit of the SILAC ratios for the unquantifiable PTM sites.

### Multiplexed staining and volumetric fluorescence imaging of cell nuclei

SDHC KO and control cells were plated at density of 1.6×10^5^ cells per 2 mL media into poly-D-lysine coated 35-mm dishes (MatTek cat# P35GC-0-10-C). For drug treatments, 10 µM C646 (Sigma Aldrich cat# SML0002) (or DMSO vehicle), 100 µM MB-3 (Sigma Aldrich cat# M2449) (or DMSO vehicle), 1 µM oligomycin A (Sigma Aldrich cat# 75351) (or DMSO vehicle) or 20 µM EX-527 (Sigma Aldrich cat# E7034) (or EtOH vehicle) was added for the specified length of time. Media was then aspirated from plates and cells were washed with 1 mL room temperature PBS. Cells were then fixed with 3.7% formaldehyde solution in PBS for 20 min at room temperature, permeabilized with PBS containing 0.1% Triton X-100 at room temperature for 15 min, and blocked with PBS containing 10% FBS for 30 min. Cells were then immunostained in blocking buffer containing 1:200 diluted mouse anti-H3K27me3 (Abcam cat# ab6002) and 1:500 diluted rabbit anti-H3K27ac (Novus Biologicals cat# NBP2-54615) with gentle agitation at room temperature for 60 min. Antibodies were then aspirated and cells were washed 3× with PBS for 5 min. Solutions of secondary antibodies were prepared in blocking buffer containing goat anti-rabbit IgG Alexa Fluor 488 (Invitrogen cat# A-11008) and goat anti-mouse IgG Alexa Fluor 594(Invitrogen cat# A32740), and added to cells for 30 min with gentle agitation, protecting from light. Cells were then washed 2× with PBS and stained with 5µg/mL DAPI in PBS for 5 min at room temperature. Cells were then washed 2× with PBS and imaged with a Zeiss LSM 780 confocal microscope using a 100× oil-immersion objective in z-stacked mode with 0.31 µm slice thickness and 0.14 µm horizontal pixel dimension. The DAPI channels from individual SDHC KO and control image stacks were used to generate a single three-state pixel classification machine learning model (extranuclear, nucleoplasm, and puncta) for performance of automated nucleus segmentation using Trainable Weka Segmentation (Arganda-Carreras et al., 2017). CellProfiler 3.0 (McQuin et al., 2018) was then used to perform automated image analysis of immunofluorescence patterns and for measurement of heterochromatin volumes.

## Supplemental item titles

**Dataset S1. Polar metabolomics data.xlsx**

**Dataset S2. ATAC-seq read mapping.xlsx**

**Movie S1. FRAP mEmerald-RAD21 control.mp4**

**Movie S2. FRAP mEmerald-RAD21 SDHC KO.mp4**

**Movie S3. FRAP mEmerald-STAG2 control.mp4**

**Movie S4. FRAP mEmerald-STAG2 SDHC KO.mp4**

